# A unique human cord blood CD8^+^CD45RA^+^CD27^+^CD161^+^ T cell subset identified by flow cytometric data analysis using Seurat

**DOI:** 10.1101/2023.08.01.549954

**Authors:** Julen Gabirel Araneta Reyes, Duan Ni, Brigitte Santner-Nanan, Gabriela Veronica Pinget, Lucie Kraftova, Thomas Myles Ashhurst, Felix Marsh-Wakefield, Claire Leana Wishart, Jian Tan, Peter Hsu, Nicholas Jonathan Cole King, Laurence Macia, Ralph Nanan

## Abstract

Advances in single cell analysis, especially cytometric approaches, have profoundly innovated immunological research. This has resulted in an expansion of high dimensional data, posing great challenges for comprehensive and unbiased analysis. Conventional manual analysis thus becomes untenable, while most computational methods lack flexibility and interoperability, hampering usability. Here, for the first time, we adapted Seurat, a single cell RNA sequencing (scRNA-seq) analysis package, for end-to-end flow cytometric data analysis. We showcased its robust analytical capacity by analyzing the adult blood and cord blood T cell profiles, which was validated by Spectre, another cytometric data analysis package, and manual analysis. Importantly, a unique CD8^+^CD45RA^+^CD27^+^CD161^+^ T cell subset, was identified in cord blood and characterized using flow cytometry and scRNA-seq analysis from a published dataset. Collectively, Seurat possesses great potential for cytometric data analysis. It facilitates thorough interpretations of high dimensional data using a single pipeline, implementing data-driven investigation in clinical immunology.

## 1. Introduction

The rapid development of analytical technologies at a single cell level over recent decades has revolutionized biological and medical research, particularly in the field of immunology. Immune cell populations are well-known for their heterogeneity and tools such as flow cytometry (or fluorescence-activated cell sorting, FACS), cytometry by time-of-flight mass spectrometry (CyTOF) and single cell RNA-sequencing (scRNA-seq), facilitate an in-depth identification and characterisation of various immune cell types [1]. Conventionally, analysis of cytometric data (including flow, spectral and mass cytometry) has relied on manual analysis based on empirical gating strategies under expert supervision. This is extremely labour-intensive and tedious, as the complex cytometric data is limited to permutational visualization of two-dimensional (2D) plots (FACS plots). These plots feature different pairs of marker combinations, which requires arduous sequential inspection [2]. The possible combinations of markers from a given panel increase exponentially with the addition of extra parameters. As more and more state-of-the-art cytometric panels exceed 20 markers [3,4], thorough manual gating analysis is becoming increasingly challenging and impractical [5]. Furthermore, such analytical workflows are inevitably subject to bias, considering their dependence on empirical knowledge and subjective selection and inspection of markers. These limitations hamper analyses and potentially conceal novel findings.

Various computational approaches have been developed as potential solutions, including methods for dimension reduction (such as t-distributed stochastic neighbour embedding, tSNE [6], and uniform manifold approximation and projection, UMAP [7]), clustering (such as PhenoGraph, and FlowSOM [8,9]), and automated cell gating and classification [5,10]. These tools all accelerate high dimensional data analysis. Moreover, they have revolutionized cytometry-based research, transitioning from the conventional hypothesis-driven strategy that focuses on specific cell types or markers, to more unbiased and comprehensive methods, that simultaneously take all data into account. Despite their notable success, these tools still suffer significant limitations. For example, many of these computational modalities are separate, and some even require specific data formatting and processing procedures. This is not user-friendly and hampers their usability and accessibility in the broader research community. While some integrative toolkits combine these modalities and offer end-to-end analysis of cross-platform cytometric data including normalisation, integration and clustering, such as Spectre [10], and ImmPort Galaxy [11], these are still few in number. There are also some commercial toolkits of this kind like OMIQ [12,13] and Cytobank [14], but they usually require paid subscriptions, limiting their availability. Furthermore, owing to their non-open-source nature, commercial toolkits can lag in flexible customizing services, as well as community-driven support, maintenance and improvements, potentially retarding optimal usability and adaptability. Hence, there is great interests in more accessible and adaptable tools.

Computational analyses of complex cytometric data have considerably benefited clinical immunological research. On one hand, clinical immunological data is notorious for its heterogeneity, highlighting the need for computational tools for data cleaning, batch alignment, and unbiased analysis. On the other hand, given the limited availability of clinical samples (such as samples with rare disease background or longitudinal samples), expanding the markers and dimensionality of cytometry panels may help to achieve more comprehensive and efficient investigation, and represents an unprecedented opportunity for high dimensional data analysis. Leveraging the rapidly developing computational analytical tools for clinical immunological studies is thus emerging as a promising avenue to provide more detailed insights into clinical contexts whilst maximising the values of limited clinical samples.

An area of increasing interest is the characterisation of immune profiles in umbilical cord blood (CB) compared to adult blood (AB). The striking immunological differences between CB and AB not only offer critical insights for disease pathogenesis, but also provides an ideal scenario for showcasing the analytical power of computational toolkits in clinical applications.

The main components of the CB immune compartment are cord blood mononuclear cells (CBMCs), well-known to exhibit unique characteristics relative to peripheral blood mononuclear cells (PBMCs) from AB, due to the semi-allogeneic environment of pregnancy. Mirroring the foetal immune system [15], CBMCs feature a more naïve phenotype [16–18], and are implicated in the physiology and pathology during both pregnancy and later in life [19–22]. Hence, understanding CB immune profiles and their differences from AB, provides precious insights into immunological development at different stages, as well as sheds light on the complex immunobiology of pregnancy.

Here, we developed a 20-marker antibody panel for thoroughly immunophenotyping T cells in both CB and AB. We adapted Seurat, a widely used end-to-end package originally for scRNA-seq analysis, for the resulting high dimensional flow cytometric data analysis. This workflow identified several previously underappreciated T cell subsets in AB, validated by Spectre, another computational cytometric analytical package, and conventional manual gating, showcasing the capacity of Seurat. Importantly, using Seurat for comparative study of CB and AB profiles, we revealed a unique CB T cell population, characterised as CD8^+^CD45RA^+^CD27^+^CD161^+^ T cells. Analysis of previously published scRNA-seq data confirmed this identified population and hinted its possible cytotoxic and pro-inflammatory properties. Together, this represents the first application example of using Seurat as a complete flow cytometric analysis workflow and demonstrated its robust analytical performance. It emerges as a simple and easy-to-use toolkit for cytometric data analysis, particularly for its pre-existing wide scRNA-seq user community. Seurat also features as a single platform but with various supplementary tools and plugins facilitating single cell analysis of both protein and RNA data, as well as their comparisons and cross-validation. This represents a novel unbiased discovery tool for complex single cell data analysis in clinical immunology.

## 2. Material and Methods

### 2.1. Subject and sample collection and storage

#### 2.1.1. Ethics approval

This study was reviewed and approved by the Human Research Ethics Committee of the Nepean Blue Mountains Local Health District according to the Declaration of Helsinki. All participants were recruited on a volunteer basis. Participants, or their guardians signed a written informed consent before sample collection.

#### 2.1.2. Adult blood (AB)

Five AB samples from healthy adult volunteers were obtained by venipuncture and collected into lithium-heparin tubes (Becton Dickinson, San Jose, CA, USA). Volunteers were excluded if they were with any intercurrent illnesses like common colds, or were on steroids, immunosuppressive drugs, or chemotherapy.

#### 2.1.3. Cord blood (CB)

Five CB samples from healthy full-term neonates born at Nepean Hospital were collected from clamped umbilical cords into lithium-heparin tubes (Becton Dickinson, San Jose, CA, USA) immediately after delivery. Women with chronic villitis, chorioamnionitis, placenta previa, gestational diabetes, or twin pregnancies were excluded.

#### 2.1.4. Mononuclear cell preparation

All blood samples were processed within 2 hours of collection. Mononuclear cells from AB and CB were isolated from whole blood through gradient centrifugation over Ficoll Paque Plus density gradient medium (Thermo Fisher Scientific, Waltham, MA, USA), washed and cryopreserved according to standard procedures [23].

### 2.2. In vitro cell culture and stimulation

Cryo-preserved PBMCs and CBMCs were thawed in a water bath at 37 °C and then washed twice with complete RPMI 1640 media (ThermoFisher Scientific) supplemented with 10% foetal bovine serum (FBS) (ThermoFisher Scientific), 2mM Glutamax and 100 U/ml Penicillin and 100μg/ml Streptamycin (ThermoFisher Scientific). Cells were then counted and checked for viability before culturing them in complete RPMI media overnight, under standard culture conditions (37°C, pH 7.4, 5% CO2, 85–95% humidity). To measure cytokines, samples were stimulated with phorbol myristate acetate (PMA) (0.1nM/ml, Sigma–Aldrich, St. Louis, MO, USA) and ionomycin (750ng/ml, Sigma Aldrich) and BD Golgi stop protein transport inhibitor, containing Monesin (Becton Dickinson) for a period of 5 hours before harvest for staining.

### 2.3. Flow Cytometric Staining

After *in vitro* culture, cells were washed with FACS buffer (phosphate-buffered saline (PBS), 0.4mM ethylenediaminetetraacetic acid (EDTA), 2%FBS) twice before staining with live/dead fixable Near IR stain (ThermoFisher Scientific) and Human TruStain FcX™ FcR block (Biolegend, San Diego, CA, USA) for 30 minutes at 4 °C. Cells were then washed twice in FACS buffer and stained with anti-CD45 antibodies for barcoding as outlined in Supplementary Table 1. Cells were incubated for 30 minutes at 4 °C and washed with FACS buffer. All 5 samples were then pooled and stained for surface markers in 70μl of the antibody cocktail (Supplementary Table 2) in BD Horizon Brilliant Stain Buffer Plus (BD Biosciences, San Jose, CA) for 30 minutes at 4 °C. Cell pellets were then washed and fixed with 100 μl BD TF Fix/Perm (intranuclear staining) or BD Cytofix Fixation buffer Solution (cytokine staining) for 20 min at room temperature in the dark. For intracellular or intranuclear marker staining, samples were then washed with 1× TF Perm/Wash buffer or Cytofix/Cytoperm buffer. Permeabilized cells were stained in 70μl of antibody cocktail (Supplementary Table 2) for 20 minutes at 4 °C. After washing, cells were resuspended in FACS buffer and analysed on the 5-laser Aurora Spectral cytometer (Cytek Biosciences, USA) on the same day.

Cytometric data was first unmixed on SpectroFlo and analysed using FlowJo (BD Life Sciences) following the gating strategy in Supplementary Fig. S1A, based on the 20-marker antibody panel described in Table 1. In brief, the panel was designed for comprehensive characterisation of T cell functional profiles, with the addition of anti-CD4 or anti-CD8 to profile CD4^+^ or CD8^+^ T cells respectively. It incorporated 15 surface markers and 5 intracellular markers in total. All analyses were carried out on the CD3^+^CD4^+^ and CD3^+^CD8^+^ T cell populations gated following the gating strategy in Supplementary Fig. S1A. For analyses with Seurat and Spectre, the CD3^+^CD4^+^ and CD3^+^CD8^+^ T cell populations were exported as CSV files with their scaled values (i.e., unmixed but not transformed values) for further analysis in RStudio.

**Table 1.** A 20-marker antibody panel for comprehensive profiling of T cells in human peripheral blood mononuclear cells (PBMCs).

| Marker Target | Staining | Markers / Functional Implications | References |
| --- | --- | --- | --- |
| **CD45** | Surface | Leukocytes (excludes erythrocytes and platelets) |  |
| **CD3** | Surface | T lymphocytes |  |
| **CD4/CD8** | Surface | Helper T cells/Cytotoxic T cells |  |
| **CD45RA** | Surface | High expression identifies naïve T cells | PMID: 18635582<br>PMID: 9348298 |
| **CD27** | Surface | Highly expressed in naïve T cells but lost in fully differentiated cells after persistent antigen stimulation | PMID: 9348298<br>PMID: 15500623 |
| **Integrin $\beta 7$ ($\beta 7$)** | Surface | Gut homing marker. | PMID: 19860663 |
| **CD161** | Surface | Type 17 responses and cytotoxic functions. | PMID: 22566826<br>PMID: 16365412 |
| **PSGL-1 (CLA)** | Surface | Skin homing marker. | PMID: 33833765 |
| **CXCR-5** | Surface | Follicular T cell marker. | PMID: 21471443 |
| **LAG-3** | Surface | Inhibitory receptor upregulated in recently activated lymphocytes. | PMID: 34067904 |
| **FcERI (FCER1A)** | Surface | High affinity receptor for IgE. | PMID: 24963147 |
| **CD49b** | Surface | Integrin ($\alpha 2\beta 1$) that binds to collagen and modulates T cell stimulation, cytokine production and survival. | PMID: 20806289<br>PMID: 17521731 |
| **CD137** | Surface | Costimulatory receptor, promoting proliferation and survival, upregulated in activated lymphocytes. | PMID: 14648705 |
| **CRTH2** | Surface | Type 2 responses. | PMID: 11168006 |
| **CD40L** | Surface | Costimulatory ligand expressed on activated T cells. | PMID: 19426221 |
| **T-box protein 21 (T-bet)** | Intracellular | Type 1 responses and effector CD8<sup>+</sup> cells differentiation | PMID: 21182077 |
| **GATA-3** | Intracellular | Type 2 responses and cytotoxic CD8<sup>+</sup> function | PMID: 23225883 |
| **BCL-6** | Intracellular | Involved in follicular helper T cell differentiation.<br>Involved in granzyme B production and memory differentiation in CD8<sup>+</sup> T cells. | PMID: 19628815<br>PMID: 31175159 |
| **Forkhead box protein P3 (FoxP3)** | Intracellular | Transcription factor expressed by regulatory T cells. | PMID: 21312192 |
| **Nur77** | Intracellular | Transcription factor indicative of T cell activation. | PMID: 30072431 |

### 2.4. High dimensional flow cytometry data analysis

#### 2.4.1. Data preparation and normalisation

For analysis using Seurat and Spectre, the exported data were first loaded in RStudio. Data then underwent an arcsinh transformation (co-factor = 2000) using Spectre (*do.asinh*). The processed data were then analysed by Spectre and Seurat.

#### 2.4.2. Analysis using Seurat

The clinical samples obtained were heterogeneous and contained differing numbers of cells. To avoid bias and skewing, the processed cytometric data was first randomly downsampled to the same number among samples before loading in Seurat for analysis. Along the conventional Seurat analysis workflow [24–27], quality check (QC) and normalisation steps originally for transcriptomic data were skipped and the data scaling process was excluded by selecting “do.scale = FALSE, do.center = FALSE” in the *ScaleData()* function.

After that, principal component analysis (PCA) was performed. Based on the PCA scores, the top PCs contributing to 99% of variance were selected for the subsequent cluster analysis using the *FindNeighbours* and *FindClusters* functions. Markers for each cluster were identified using the *FindMarkers()* function based on the default settings.

#### 2.4.3. Analysis using Spectre

For analysis with Spectre, input data were first downsampled and then analysed following Ashhurst et al, 2022 [10], except a 5×5 self-organizing map (SOM) was used. The final clustering numbers were also adjusted to the same as Seurat’s results for comparison.

### 2.5. Single cell RNA sequencing (scRNA-seq) analysis

Single cell RNA sequencing (scRNA-seq) data was downloaded from Gene Expression Omnibus from a previous study [28] (GEO: GSE158493). Processed data after normalisation and scaling are available from the original manuscript. Since the original dataset was grouped based on sample origins (foetal spleen, full-term umbilical cord blood and adult peripheral blood), the LIGER package was used to re-integrate the data with different origins to eliminate potential batch effects [29]. After that, the data were analysed and visualized with Seurat 4.3.0. [24–27]. Marker genes were identified using Seurat’s *FindMarkers* function based on its default settings, with thresholds for differentially expressed genes defined as p < 0.05 and LOG2(Fold change) higher than 0.5 or lower than -0.5. Gene set enrichment analysis (GSEA) was run with package fgsea and ClusterProfiler following their tutorials [30,31].

### 2.6. Statistics

Statistical analysis was performed with PRISM (GraphPad) or in RStudio (v4.1.2). Mann-Whitney test was used for comparison between two groups. Differences were considered to be statistically significant when p < 0.05.

## 3. Results

### 3.1. Seurat is a reliable tool for high dimensional flow cytometry data analysis

Analysis of human peripheral blood, and the heterogenous T cell compartments therein, is of great importance considering the availability of blood samples and its ubiquitous use in medical research and clinical diagnosis. Here, we established a 20-marker antibody panel (Table 1) to comprehensively characterise the functional profiles of CD4^+^ and CD8^+^ T cells in human peripheral blood mononuclear cells (PBMCs). Details about the markers and their functions are described in Table 1. They cover various aspects of T cell biology, including maturation, activation, migration, and function.

PBMCs from healthy adult individuals were collected and processed based on the workflow in Fig. 1A. To reduce antibody wastage as well as minimize the inter-sample variations caused by batch effects introduced during experimental processes, which is an intrinsic problem for clinical studies involving flow cytometry, we utilised a barcoding system leveraging the anti-CD45 labelling. In the current study, PBMC samples were stained with our 20-colour antibody panel and analysed by spectral cytometry. After demultiplexing, the resulting cytometric data were manually gated for CD4^+^ and CD8^+^ T cell populations (Supplementary Fig. S1A).

**Fig. 1.**
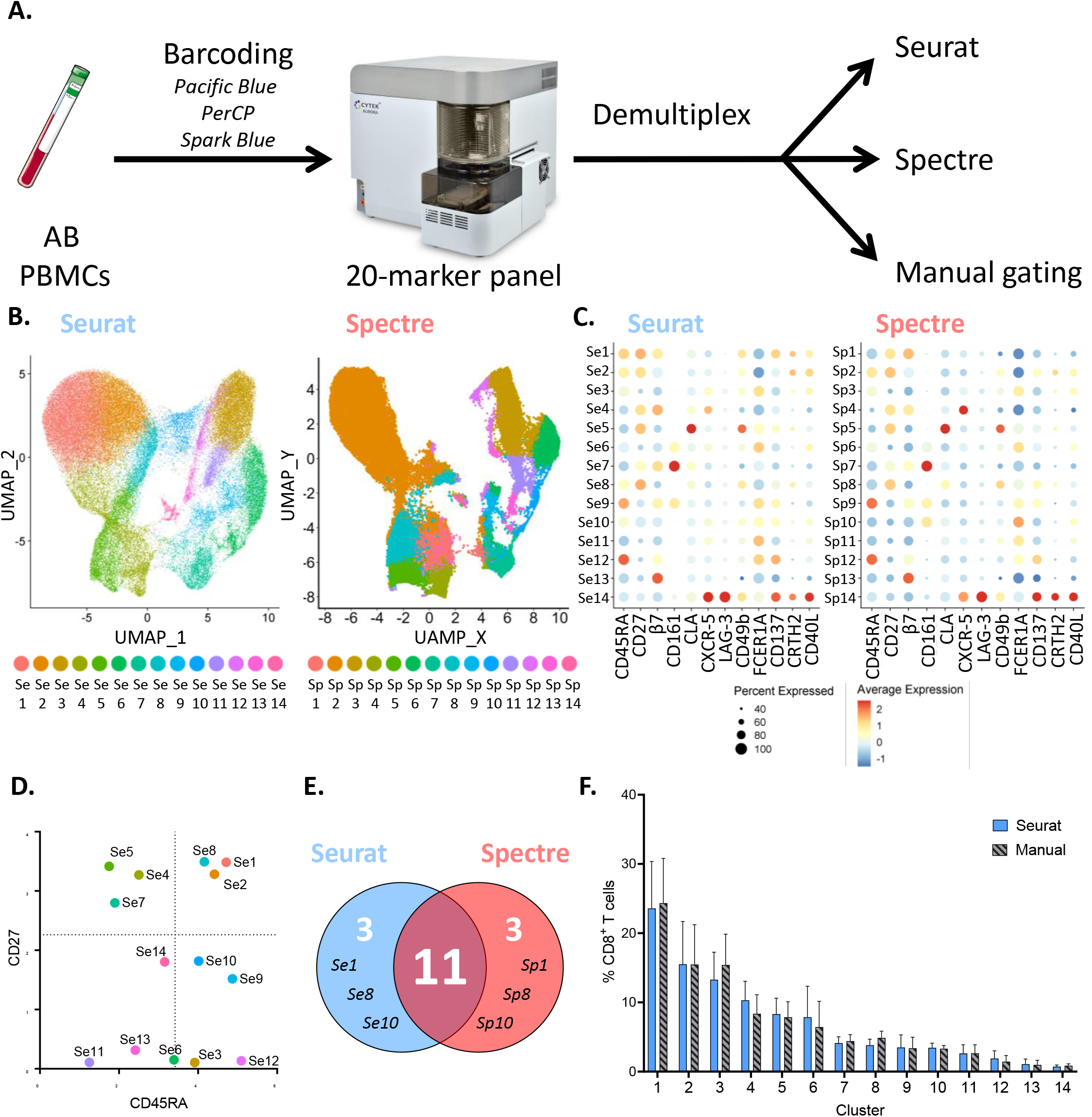
Adapting Seurat for high dimensional flow cytometric data analysis retrieved robust results on adult blood (AB) peripheral blood mononuclear cells (PBMCs), confirmed by Spectre and manual analysis. **A.** An overview of the adult blood (AB) study design. Peripheral blood mononuclear cells (PBMCs) were isolated from blood samples from healthy adult and then were first labelled with anti-CD45 antibodies with different fluorophores or their combinations for barcoding. After that, PBMCs were pooled together and stained with our newly developed 20-marker antibody panel and analysed with Spectral Cytometry. Next, the resulting data were first demultiplexed based on their CD45 marker signals and then subject to analysis with Seurat, Spectre, and manual gating. **B.** Uniform manifold approximation and projection (UMAP) plots visualizing the clustering results from Seurat (left) and Spectre (right) based on the adult PBMC CD8^+^ T cell experiment. One colour represents one cluster. **C.** Dot plots visualizing the clusters identified by Seurat (left) and Spectre (right) and their marker expression profiles. The size of the dot corresponds to the percentage of cells expressing the corresponding markers and the colour gradient reflects the average normalised expression of the corresponding markers. **D.** Projection of 14 clusters identified by Seurat based on the adult PBMC CD8^+^ T cell experiment onto the two-dimensional (2D) plot comparing their expression of CD27 and CD45RA. The dashed lines denote the average normalised expression of CD27 and CD45RA for all cells. **E.** Venn diagram comparing the clustering results from Seurat and Spectre. Both methods were set to generate 14 clusters and 11 out of 14 clusters could be identified by both methods, while Se1, Se8, and Se10 could only be identified by Seurat and Sp1, Sp8 and Sp10 could only be identified by Spectre. **F.** Bar chart comparing the proportions per sample within the total CD8^+^ T cell compartments of the clusters identified by Seurat or retrieved by manual gating. N=5 per group and data are presented as mean ± s.e.m.

For proof-of-concept, the CD8^+^ T cell data was analysed with Seurat and compared with Spectre, based on the 12 functional markers out of the total 15 surface markers, excluding lineage markers such as CD45, CD3 and CD8. The results were validated through manual gating as well.

Seurat clustered the adult CD8^+^ T cell compartment into 14 clusters (Se1-14) and projected them onto the 2D UMAP plot as shown in Fig. 1B. Historically, T cells are gated into four populations based on expression of CD45RA and CD27. The CD45RA^+^CD27^+^ population is defined as naïve, the CD45RA^-^CD27^+^ population as central memory (CM), the CD45RA^-^CD27^-^ population as effector memory (EM) and the CD45RA^+^CD27^-^ is effector memory that re-express CD45RA (EMRA) [32]. Comparing Seurat’s results to this classification, we found that Se1, Se2 and Se8 were naïve, Se4, Se5 and Se7 were CM, Se11 and Se13 were EM, and Se12 was EMRA. Interestingly, Seurat clustering retrieved populations exhibiting intermediate expression levels of CD45RA and CD27, such as clusters Se3, Se6, Se9, Se10 and Se14 (Fig. 1C and 1D), which did not necessarily fall into any of the four conventional gates mentioned above. This implies that the conventional gating strategy based only on positive or negative expression of markers is incomplete. Seurat, on the other hand, can provide a more comprehensive analysis for multiple markers, and thus enable the discovery of previously unidentified subsets.

Feature markers identified by Seurat and the potential functional properties of each cluster are summarised in Supplementary Table 3. Clusters with naïve phenotypes (Se1, Se2 and Se8) expressed different levels of integrin β7, indicating differential gut homing potentials [33]. Two CM clusters, Se4 and Se7, were also high in integrin β7, however they could be separated based on CXCR-5 and CD161 expression, as markers for follicular T cells and cytotoxic T cells, respectively. The CM cluster Se5 uniquely featured the skin homing marker, CLA. Cluster Se6 and Se9 both expressed intermediate levels of CD161, but Se6 also had intermediate expression of integrin β7 while Se9 did not express integrin β7 at all. Se3 exhibited a similar profile to Se6, except for the absence of CD161. Additionally, their intermediate expression of CD45RA implied that they might represent the transitional state between EM and EMRA [34]. For EM-like and EMRA-like clusters, Se13 and Se12, but not Se11, they expressed high levels of integrin β7. Finally, Se14 was characterised by their high level of LAG-3, a marker for early T cell activation, while Se10 might be a transitional population during the T cell activation and maturation, considering its intermediate expression of both CD45RA and CD27. Other markers in our panel including CD49b and FCER1A were expressed at relatively low levels in T cells in the absence of stimulation. These subtle differences could still be detected by Seurat and confirmed with manual analysis (Fig. 1C and Supplementary Fig. S2A-B). For example, CD49b, a collagen-binding integrin, showed the highest level in the skin homing population Se5, while FCER1A expression inversely correlated with CD27 expression, as exemplified in Se3, Se6, Se9, Se11 and Se12. Considering their relatively low level of expressions, further research is required to evaluate whether such mild differences are of biological significance.

In parallel, we applied Spectre’s workflow to analyse our dataset, manually defining 14 clusters as the final clustering output to cross-validate the findings from Seurat. Comparison of the Seurat and Spectre results revealed that their outcomes were similar, with 11 of the 14 clusters being identified by both approaches, displaying comparable cluster sizes and feature marker expression (Fig. 1C, 1E and Supplementary Fig. S2C). This indicated the robust and reliable performance of Seurat. As for their discrepancies, the Sp2 identified by Spectre was divided into three distinct clusters by Seurat, Se1, Se2 and Se8 (Supplementary Fig. S2D). In contrast, Se4 in Seurat analysis was split into Sp1 and Sp4 by Spectre (Supplementary Fig. S2D). However, since Seurat and Spectre adopt different algorithms for their analyses, such differences were potentially to be expected.

Importantly, we further validated the results from Seurat with manual gating. As shown in Supplementary Fig. 3, all 14 Seurat clusters could be gated out according to their marker expression on 2D FACS plots. Furthermore, the proportion of each population in the total CD8^+^ T cells were also comparable between manual gating and Seurat clustering (Fig. 1F).

Finally, we similarly applied the workflow to the gated CD4^+^ T cells and retrieved 15 clusters with Seurat (Supplementary Fig. S4A-B). These subsets could also be similarly confirmed by manual gating and cross-validated with Spectre, which identified 12 out of the 15 subsets obtained by Seurat (Supplementary Fig. S4C-D).

Together, these results demonstrated that Seurat, a tool originally developed for scRNA-seq data analysis, is also applicable and robust for high-dimensional flow cytometric data analysis. Its analysis helps to characterize the CD8^+^ T cells in PBMCs in more details, retrieving novel T cell sub-clusters.

### 3.2. Comparative profiling of CD8^+^ T cells in adult blood and cord blood

Human circulatory T cells are plastic across the human lifespan and might be linked to the differential disease susceptibility across human lives [35]. We next aimed to compare the profiles of the T cell compartments from the cord blood (CB) and adult blood (AB) using Seurat.

As shown in Fig. 2A, peripheral blood mononuclear cells (PBMCs) from AB and cord blood mononuclear cells (CBMCs) from CB were analysed based on our 20-parameter antibody panel first with manual gating. Next, the high dimensional data were analysed by Seurat and Spectre.

**Fig. 2.**
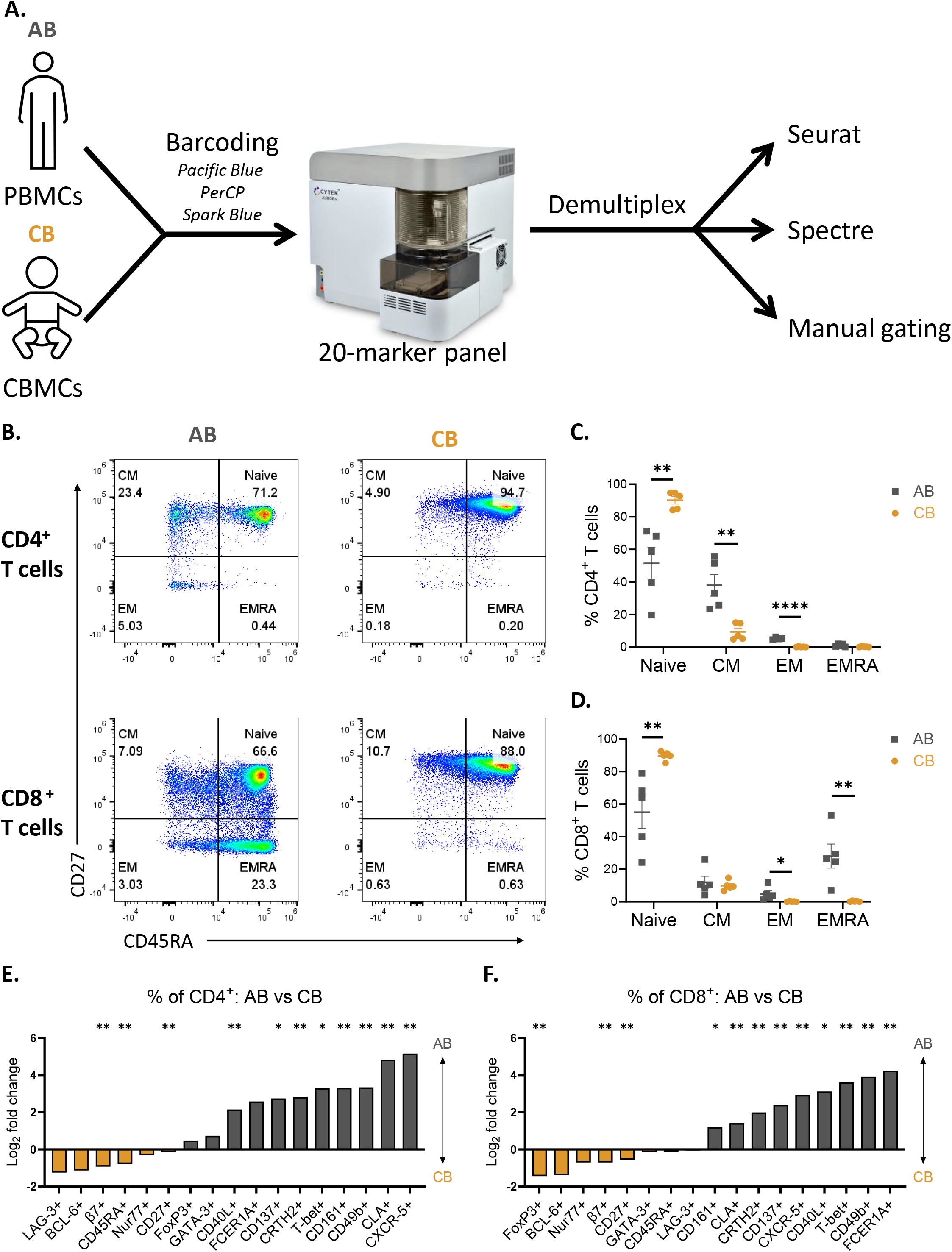
Comparative analysis of T cell profiles in peripheral blood mononuclear cells (PBMCs) from adult blood (AB) and cord blood mononuclear cells (CBMCs) from cord blood (CB) **A.** An overview of the adult blood (AB) and cord blood (CB) study design. Peripheral blood mononuclear cells (PBMCs) and cord blood mononuclear cells (CBMCs) were isolated and then were first labelled with anti-CD45 antibodies with different fluorophores or their combinations for barcoding. After that, PBMCs were pooled together and stained with our newly developed 20-marker antibody panel and analysed with Spectral Cytometry. Next, the resulting data were first demultiplexed based on their CD45 marker signals and then subject to analysis with Seurat and manual gating. **B.** Representative flow cytometric plots of CD4^+^ and CD8^+^ T cells from AB and CB to identify naïve (CD45RA^+^CD27^+^), central memory (CM, CD45RA^-^CD27^+^), effector memory (EM, CD45RA^-^CD27^-^) and effector memory cells re-expressing CD45RA (EMRA, CD45RA^+^CD27^-^) subsets. **(C-D)** Scatter bar charts for the proportions of naïve, CM, EM, and EMRA subsets within CD4^+^ **(C)** and CD8^+^ **(D)** T cells from AB and CB. **(E-F)** Bar charts for the log2(fold change) comparing AB versus CB for the proportions of populations expressing the corresponding markers among CD4^+^ (E) and CD8^+^ (F) T cells. The asterisks denote the populations whose proportions are significantly different between AB and CB. N=5 per group and data are presented as mean, with * p<0.05 and ** p<0.01 by unpaired Mann-Whitney t-test.

The proportion of CD4^+^ and CD8^+^ T cells among total lymphocytes showed no difference in PBMCs and CBMCs (Supplementary Fig. S1B). As previously shown [16], there were more naïve T cells and fewer EM T cells in both CD4^+^ and CD8^+^ compartments from CBMCs compared with PBMCs (Fig. 2B-D). Reflective of a mature phenotype, adult PBMCs had higher proportions of CD4^+^ CM T cells and CD8^+^ EMRA T cells (Fig. 2C-D). Our comprehensive 20-parameter antibody panel enabled an in-depth characterisation of the functional status of T cells.

As shown in Fig. 2E, CD4^+^ T cells in adult PBMCs had higher proportions of cells expressing T-bet, CLA and CXCR-5, while more CD4^+^ T cells from CB expressed CD27, β7, and LAG-3. Similarly, CB CD8^+^ T cells had a higher expression of CD27, β7, alongside FoxP3, whilst adult CD8^+^ T cells had higher proportions of cells expressing CD161, CLA, CXCR-5 and T-bet (Fig. 2F and Supplementary Fig. S5A-B). These findings are consistent with previous reports of the higher gut homing potential of CBMCs, whilst PBMCs are more likely to migrate to the skin [16,35–37]. Moreover, the lower expression of T-bet and overall naïve-biased phenotype of CB T cells coincides with their reduced IFN-γ, IL-4 and IL-13 production compared to AB T cells (Supplementary Fig. S5C-D) [38].

### 3.3. Analysis of adult blood and cord blood CD8^+^ T cells with Seurat identifies a unique cord blood CD8^+^CD45RA^+^CD27^+^CD161^+^ T cell subset

Conventional manual gating workflows are empirical and subject to bias. Such analysis is less likely to potentially reveal new cell populations. Focusing on the CD8^+^ T cell compartment, we therefore performed unsupervised principal component analysis (PCA) on the AB and CB combined dataset based on expression levels quantified by median fluorescence intensity (MFI) of the 12 surface markers used for clustering (Fig. 3A). As expected, PBMC samples were distinctly separated from CBMC samples along the first PC (PC1, accounting for 18.0% of the variance), which was consistent with the differential expression of various T cell functional markers in AB samples relative to CB samples (Fig. 2E-F).

**Fig. 3.**
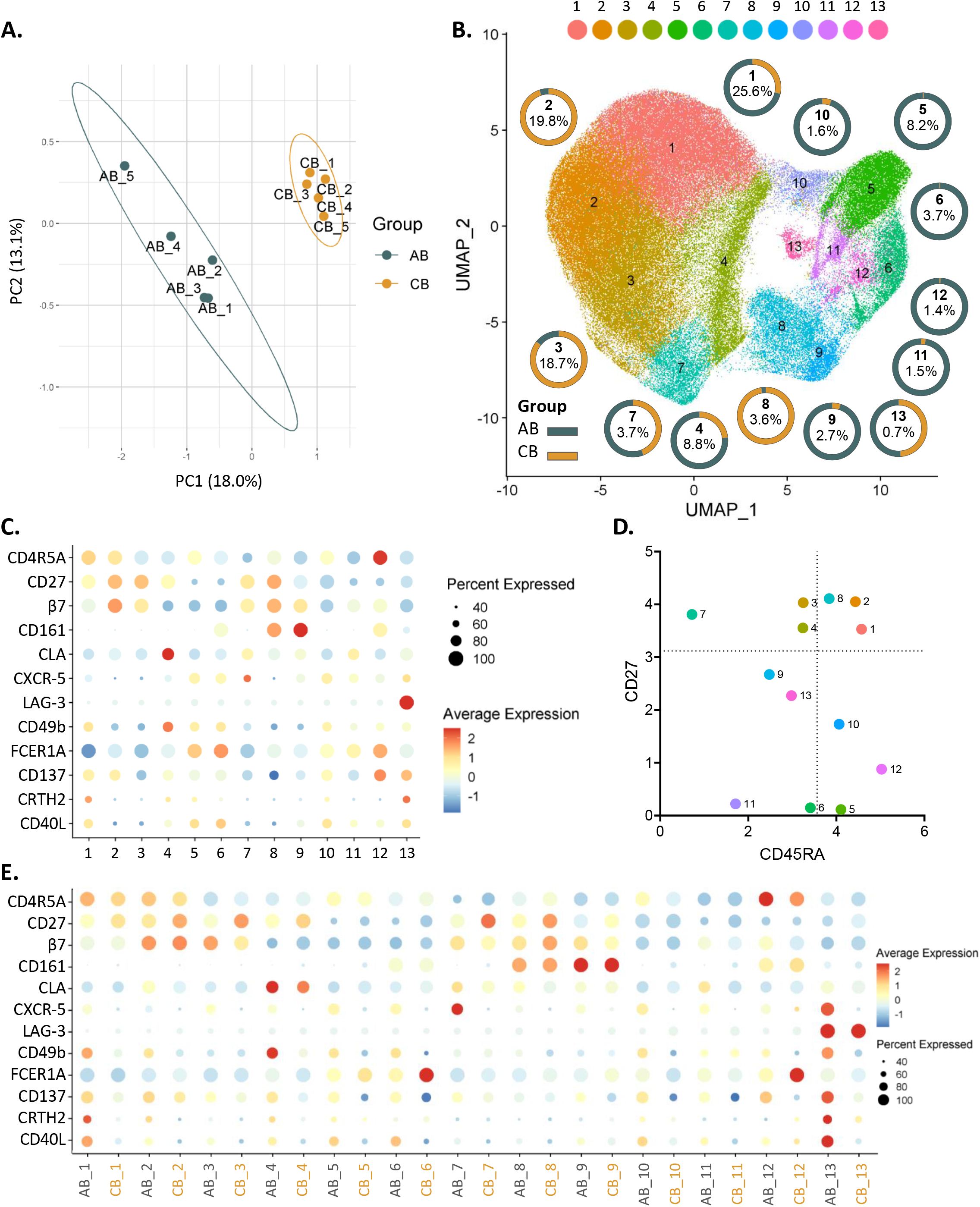
Unbiased comparative analysis of CD8^+^ T cell profiles in peripheral blood mononuclear cells (PBMCs) from adult blood (AB) and cord blood mononuclear cells (CBMCs) from cord blood (CB) with principal component analysis (PCA) and Seurat. **A.** PCA based on the marker expression levels on CD8^+^ T cells from AB and CB samples. **B.** Uniform manifold approximation and projection (UMAP) plot visualizing the clustering results from Seurat and the compositional contribution of each cluster from AB versus CB. The donut charts visualized the proportions of each cluster that are from AB (dark blue) and CB (yellow), and the numbers within denoted the proportions of the corresponding clusters within the overall CD8^+^ T cells from both AB and CB. **C.** Dot plot visualizing the clusters identified by Seurat and their marker expression profiles. The size of the dot corresponds to the percentage of cells expressing the corresponding markers and the colour gradient reflects the average normalised expression of the corresponding markers. **D.** Projection of 13 clusters identified by Seurat based on the AB versus CB experiment onto the two-dimensional (2D) plot comparing their expression of CD27 and CD45RA. The dashed lines denote the average normalised expression of CD27 and CD45RA for all cells. **E.** Dot plot comparing the clusters from AB versus CB identified by Seurat and their marker expression profiles. The size of the dot corresponds to the percentage of cells expressing the corresponding markers and the colour gradient reflects the average normalised expression of the corresponding markers.

Next, Seurat was used for more in-depth analyses, identifying 13 sub-populations from the combined AB and CB dataset (Fig. 3B). Seurat clustering distinguished three naïve (clusters 1, 2 and 8), three CM (clusters 3, 4 and 7), one EM (cluster 11), and two EMRA clusters (clusters 5 and 12), with the remaining clusters exhibiting intermediate expression levels of CD45RA and/or CD27 (clusters 6, 9, 10, 13) (Fig. 3C-D). Details about feature markers and potential functional properties of the identified CD8^+^ T cell subsets are summarised in Supplementary Table 4. Clusters 1 and 2 shared a naïve phenotype but differed in their expression of integrin β7 and gut homing potential, consistent with previous findings demonstrating an increased expression of gut homing receptors in naïve CB lymphocytes [39]. The naïve cluster 8 was also high in integrin β7 but additionally expressed CD161, suggesting a cytotoxic phenotype. Out of the three CM subsets identified by Seurat, Clusters 3 and 7 of the CM class both had high integrin β7 expression, indicative of a migratory preference to the gut. Cluster 7 differed from Cluster 3 through its low expression of CD45RA and high expression of the follicular T cell marker, CXCR-5. Opposed to gut homing, the CM cluster 4 did not express integrin β7 but instead highly expressed the skin homing marker CLA and integrin CD49b. Integrin β7 levels could also distinguish the EMRA-like clusters 5 versus 12 as well as clusters 6 versus 9, with clusters 6 and 9 also expressing CD161. Finally, cluster 13 featured high levels of LAG-3 and CXCR-5.

We again compared the analyses from Seurat and Spectre based on the combined AB and CB dataset. Both methods identified 13 clusters, of which 11 were commonly shared, with comparable feature markers (Supplementary Fig. S5E-F). Similarly, Seurat clusters in both AB and CB could also be validated through manual gating (Supplementary Fig. S6 and S7). These results again highlighted the robust performance of Seurat for analysing high dimensional cytometric data.

Striking differences in the abundance of the Seurat clusters were found comparing AB and CB samples (Fig. 3B and Supplementary Fig. S8A). The naïve cluster 1 was predominantly abundant in AB whilst cluster 2 consisted primarily of CB cells. Seurat cluster analysis was able to separate these two naïve populations based on the differential expression of integrin β7 and CD27, which were enriched in the naïve population from CB (cluster 2) [16,39]. Consistent with previous studies comparing adult and cord blood, CD8^+^ effector memory populations were almost exclusively found in AB [40]. This includes all the identified EM (cluster 11) and EMRA (clusters 5 and 12) subsets, in addition to clusters 6, 9 and 10. Unlike these EM subsets, CM populations were present in both AB and CB. Of note, cluster 3 was dominant in CB whilst clusters 4 and 7 were enriched in AB. Finally, the LAG-3^+^ cluster 13 was equally abundant in both AB and CB.

Moreover, we also documented differential expression of various markers between the clusters from AB and CB. Reflective of their naïve phenotype and lack of antigenic exposure, several CB CD8^+^ T cell populations (clusters 1, 2, 3, 4, 7, 8, 9) exhibit higher expression of CD27 compared to their adult equivalents (Fig. 3E). In contrast, the expression of CXCR-5 in clusters 7 and 13 alongside CLA expression in cluster 4 was higher in adult CD8^+^ T cells compared to CB (Fig. 3E).

Intriguingly, our clustering analysis identified a population (cluster 8) that is almost exclusive to CBMCs (7.11% of CD8^+^ T cells in CB versus 0.19% in AB) (Fig. 4A-C). This cluster is characterised as CD8^+^CD45RA^+^CD27^+^CD161^+^, partly overlapping with a previously reported but not fully characterised, CD8^+^CD161^+^ T cell population found in CB [41,42]. Analysis based on our 20-parameter panel provided an unprecedented functional overview of the CD8^+^CD45RA^+^CD27^+^CD161^+^ T cell subset. We discovered that this newly identified population had high integrin β7 expression (Fig. 4B and Fig. 4D) as well as higher CLA, BCL-6, T-bet, and GATA-3, but lower FoxP3 levels compared to its CD161^-^ counterpart (Fig. 4E). No differences in LAG-3, CXCR-5, Nur77, CD137 and FCER1A were found between the CD8^+^CD45RA^+^CD27^+^CD161^+^ and CD8^+^CD45RA^+^CD27^+^CD161^-^ T cells (Fig. 4E). Upon stimulation with PMA and ionomycin, this subset predominantly produced IFN-γ and IL-4 but lowly expressed IL-5, IL-10, and IL-13. It only differed significantly in IL-10 production relative to the CD161^-^ counterpart (Fig. 4F). Based on these results, the CD8^+^CD45RA^+^CD27^+^CD161^+^ sub-population (cluster 8) appeared to be a pro-inflammatory and cytotoxic T cell subset.

**Fig. 4.**
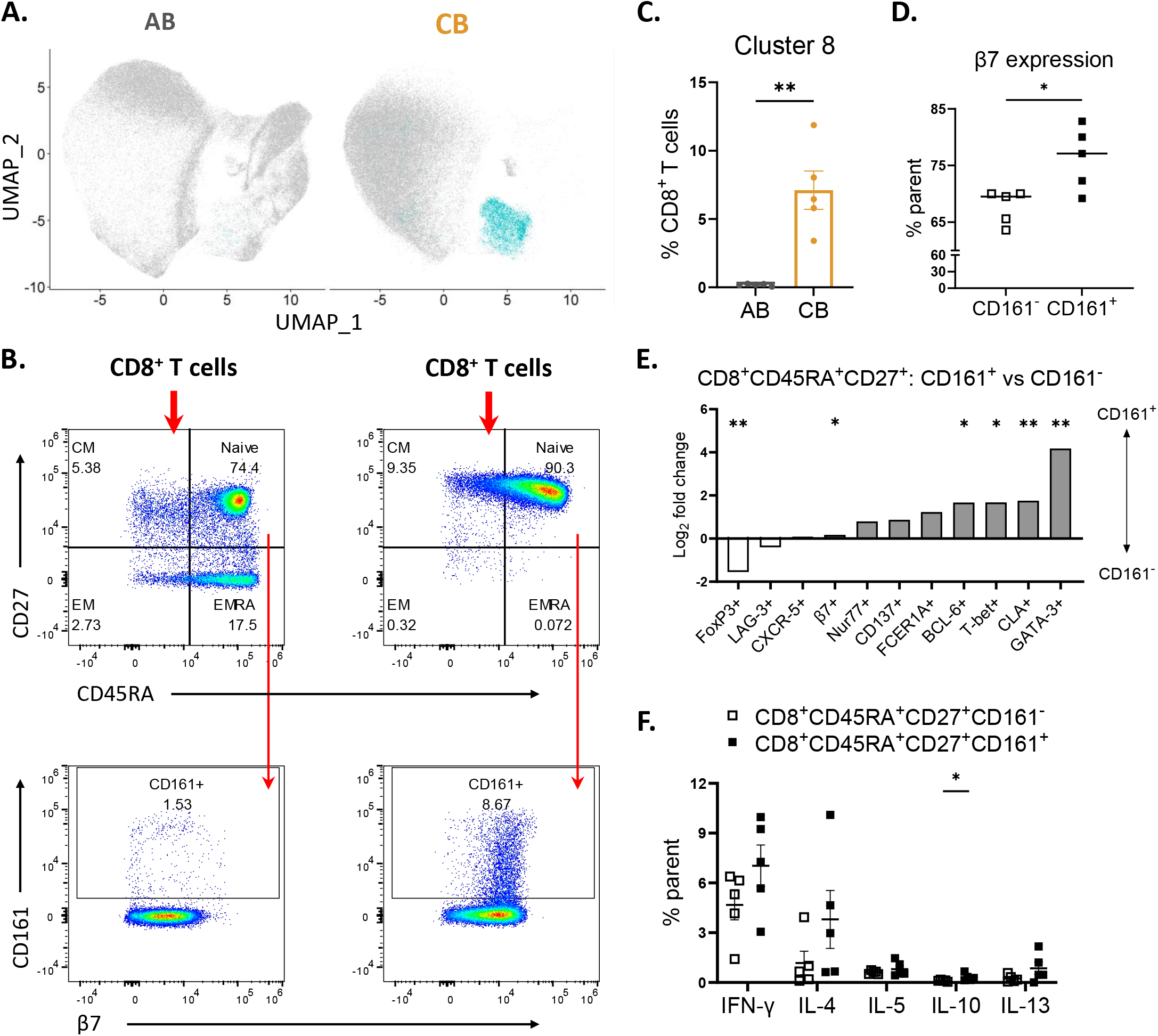
Seurat analysis identified a unique CD8^+^CD45RA^+^CD27^+^CD161^+^ T cell population in cord blood. **A.** Overlay of the newly identified CD8^+^CD45RA^+^CD27^+^CD161^+^ T cell subset (cluster 8) onto the Uniform manifold approximation and projection (UMAP) plots for AB (left) and CB (right) CD8^+^ T cell compartments. **B.** Manual gating strategies to identify the CD8^+^CD45RA^+^CD27^+^CD161^+^ T cell subset (cluster 8) from AB (left) and CB (right) CD8^+^ T cell compartments. **C**. Scatter bar chart for the proportions of CD8^+^CD45RA^+^CD27^+^CD161^+^ T cell subset (cluster 8) in AB (black) and CB (yellow) CD8^+^ T cell compartments. **D.** Scatter dot plot of the proportions of cells expression integrin β7 in CD8^+^CD45RA^+^CD27^+^CD161^+^ and CD8^+^CD45RA^+^CD27^+^CD161^-^ T cell subsets. **E.** Bar chart for the log2(fold change) comparing the proportions of populations expressing the corresponding markers among CD8^+^CD45RA^+^CD27^+^CD161^+^ and CD8^+^CD45RA^+^CD27^+^CD161^-^ T cell subsets. The asterisks denote the populations whose proportions are significantly different between AB and CB. N=5 per group and data are presented as mean, with * p<0.05 and ** p<0.01 by unpaired Mann-Whitney t-test. **F.** Scatter dot plot of the proportions of cells expressing IFN-γ, IL-4, IL-5, IL-10, and IL-13 in CD8^+^CD45RA^+^CD27^+^CD161^+^ and CD8^+^CD45RA^+^CD27^+^CD161^-^ T cell subsets.

In summary, using our high-dimensional antibody panel and Seurat analysis, we thoroughly profiled the CD8^+^ T cell compartment in AB versus CB. This revealed a unique CD8^+^CD45RA^+^CD27^+^CD161^+^ T cell subset in CB which we characterized.

### 3.4. Cross-validation and further characterisation of the CD8^+^CD45RA^+^CD27^+^CD161^+^ T cell subset using scRNA-seq

To validate our findings and further characterise the population identified by Seurat in CBMCs, we leveraged a recently published scRNA-seq dataset for naïve CD8^+^ T cells, which analysed 18513 cells across different developmental stages and compartments. This dataset covers naïve T cells from foetal spleen, umbilical cord blood and adult peripheral blood. As shown in Fig. 5A and Supplementary Fig. 9A, the overall naïve CD8^+^ T cell population (sorted as CD8^+^CD45RA^+^CD27^+^CCR7^+^CD95^-^) was further clustered into four subsets. Cluster0 expressed high levels of *RGS1*, which is linked to T cell exhaustion [43]; cluster1 was high in *IL7R* and *SELL*, indicative of a naïve phenotype; and cluster2 was marked by, which might be potentially linked to a memory T cell phenotype [44]. Intriguingly, cluster3 exhibited similar features to the population we identified, characterised by its expression of *PTPRC* (gene encoding CD45RA), *KLRB1* (gene encoding CD161), *ITG*Β*7* (gene encoding integrin β7) and *CD27* (Fig. 5B). Differentially expressed gene analysis with DESeq2 [45] found 93 genes upregulated in cluster3 compared with the remaining naïve CD8^+^ T cells, while 2 genes were downregulated (Fig. 5C). Among the upregulated genes in cluter3, there were several related to cytotoxic T cell features, such as *GZMA*, *GZMK*, *GZMM*, *CST7*, and *NKG7* [46], indicating their cytotoxic functions, which is similar to our cytometric results and the previous reports for CD8^+^ CD161^+^ T cells [41]. Additional inflammation-related genes were upregulated, such as *ID1* [47] and *CMC1* [48]. Interestingly, *CCL5*, related to a memory phenotype, was also significantly upregulated in cluster3. Additionally, several chemokine receptors were also increased, such as *CXCR3* and *CXCR4*.

**Fig. 5.**
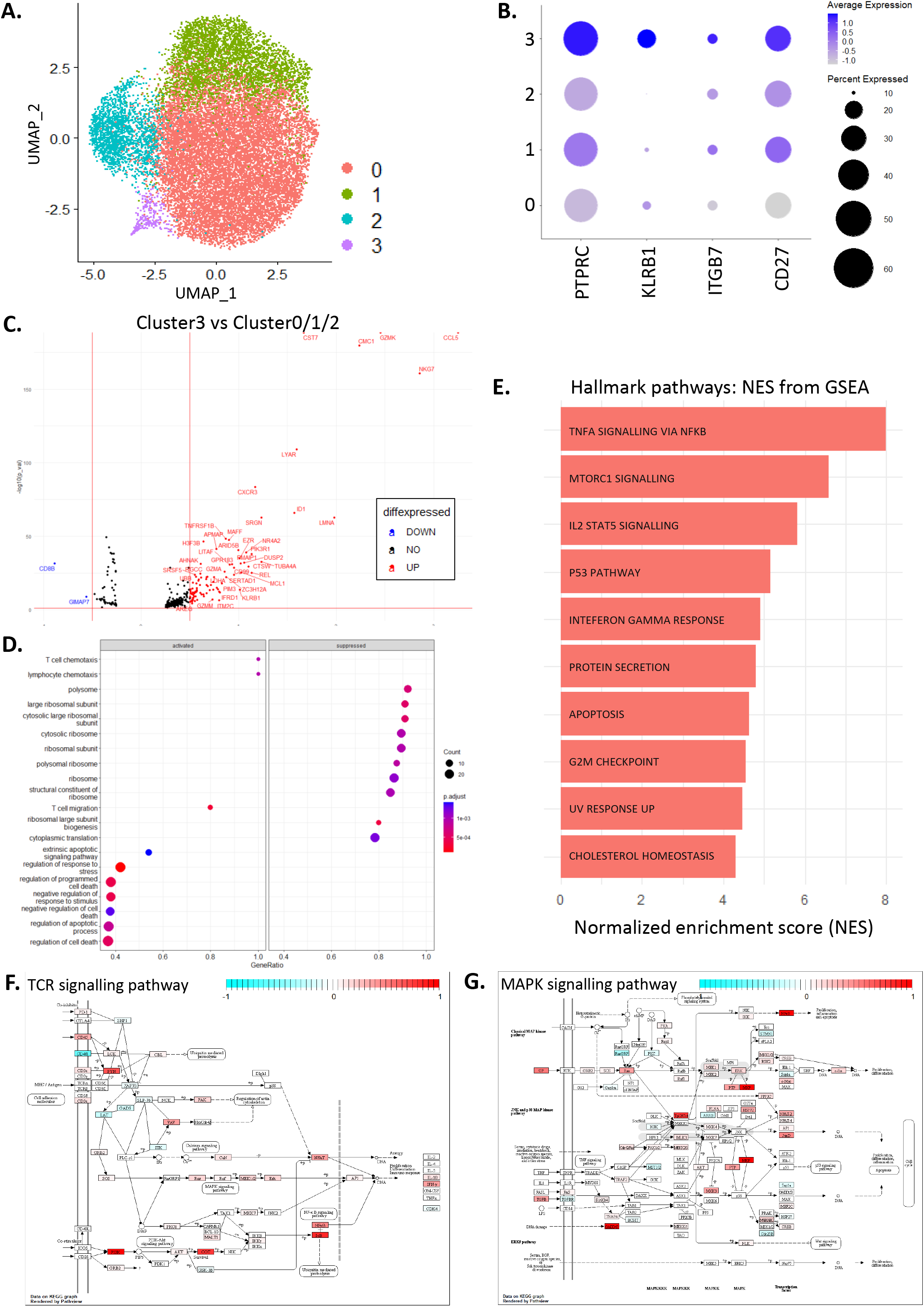
Single cell RNA sequencing (scRNA-seq) cross-validation and characterisation of the newly identified CD8^+^CD45RA^+^CD27^+^CD161^+^ T cell population. **A.** Uniform manifold approximation and projection (UMAP) plot visualizing the clustering results from re-analysis of the scRNA-seq data for naïve CD8^+^ T cells from GEO: GSE158493. **B.** Dot plot visualizing the clusters identified by Seurat from scRNA-seq analysis and their expression levels of *PTPRC, KLRB1, ITGB7* and *CD27*. The size of the dot corresponds to the percentage of cells expressing the corresponding markers and the colour gradient reflects the average normalised expression of the corresponding markers. **C.** Volcano plot showing the differentially expressed genes comparing CD8^+^CD45RA^+^CD27^+^CD161^+^ and CD8^+^CD45RA^+^CD27^+^CD161^-^ T cell subsets. The red dots indicate genes up-regulated in CD8^+^CD45RA^+^CD27^+^CD161^+^ T cells, the blue dots indicate genes down-regulated in CD8^+^CD45RA^+^CD27^+^CD161^+^ T cells, and the black ones are genes without significant changes. **D.** Dot plot for the activated (left) and suppressed (right) pathways in CD8^+^CD45RA^+^CD27^+^CD161^+^ T cell subset compared to CD8^+^CD45RA^+^CD27^+^CD161^-^ T cell subset analysed by ClusterProfiler. The size of the dot corresponds to the number of genes within the corresponding pathway, the GeneRatio is calculated by the ratio between the number of genes that are significantly differentially expressed comparing CD8^+^CD45RA^+^CD27^+^CD161^+^ and CD8^+^CD45RA^+^CD27^+^CD161^-^ T cell subsets and the number of total genes involved in the corresponding pathway, and the colour gradient reflects the adjusted p value for the comparison. **E.** The top enriched gene sets of gene set enrichment analysis (GSEA) comparing CD8^+^CD45RA^+^CD27^+^CD161^+^ and CD8^+^CD45RA^+^CD27^+^CD161^-^ T cell subsets based on the Hallmark gene sets. (**F-G**). Visualization of differentially expressed genes comparing CD8^+^CD45RA^+^CD27^+^CD161^+^ T cell subsets relative to CD8^+^CD45RA^+^CD27^+^CD161^-^ T cell subsets that are involved in T cell receptor (TCR) signalling pathway (**F**) and mitogen-activated protein kinase (MAPK) signalling pathway (**G**).

Gene Set Enrichment Analysis (GSEA) was next carried out with the differentially expressed genes using ClusterProfiler [30,31] based on the Gene Ontology pathways [49]. As shown in Fig. 5D, compared with other naïve CD8^+^ T cells, cluster3 was enriched for pathways related to *T cell chemotaxis*, *lymphocyte chemotaxis* and *T cell migration*, consistent with the differentially expressed gene analysis (Fig. 5C) and our flow cytometric data (Fig. 3I and 3J). On the other hand, this population was suppressed in pathways related to ribosomes.

These analyses were based only on differentially expressed genes, and we next conducted GSEA based on the overall transcriptomic data using the fgsea package [50]. GSEA using the *Hallmark gene set* showed that cluster3 displayed enrichment in inflammatory pathways such as *TNF*α *signalling via NF*κ*B, IL2-STAT5 signalling,* and *IFN*γ *response* (Fig. 5E). Additionally, they also showed enrichment in the *MAPK signalling pathway* and the *T cell receptor signalling pathway* (Fig. 5F, G).

In summary, scRNA-seq analysis from an independent dataset cross-validated the CD8^+^CD45RA^+^CD27^+^CD161^+^ population identified by our Seurat-based analysis. It also characterised it as a naïve subset but with a potential pro-inflammatory and cytotoxic profile.

## 4. Discussion

There has been a significant expansion of both the size and complexity of cytometric data, especially in the field of clinical immunology. Such high dimensional and complicated datasets cause great difficulties for conventional manual analytical strategies, inevitably hampering comprehensive and unbiased analyses and interpretation. Consequently, myriad computational toolkits have been developed, aiming to address these challenges, but their effective applications are sometimes restricted, suffering from a lack of flexibility and interoperability. Recently, packages such as Spectre [10] and tidyof [51] were developed, attempting to provide integrative, end-to-end services for cytometric data analysis. However, they have only been sporadically applied, due to them being standalone pipelines, requiring adaptation to completely new packages, and demanding significant prerequisite coding knowledge.

In the present study, we repurposed Seurat, a well-established package for scRNA-seq data analysis, for high dimensional flow cytometric data analysis [24–27]. Seurat has long shared great popularity within the field of single cell analysis. It is community-driven and well-supported and has more than 20 R packages for related data processing and analysis. Therefore, it is likely to be more accessible and easier to use, particularly for the broad users with previous experiences in scRNA-seq looking to flow cytometry to complement their investigative breadth.

Here, we showcased the robust capacity of Seurat, based on our experiments profiling the T cell compartments in adult blood (AB) and cord blood (CB). Overall, Seurat generated similar results to Spectre, which were also confirmed by manual gating. Importantly, with our approach, we identified a unique T cell subset (CD8^+^CD45RA^+^CD27^+^CD161^+^) within CB and cross-validated its functional profiles with an independent scRNA-seq dataset using Seurat. Together, these data highlight the great potential of Seurat for cytometric data analysis. It represents a simple single platform for the unbiased analysis of both protein and RNA data at single cell resolution. This will enable simpler comparison and cross-validation of cytometric and scRNA-seq studies and facilitate more comprehensive investigations and discoveries in clinical immunology.

A plethora of state-of-the-art mathematical algorithms or statistical models are used in the field of cytometry computational analysis. These include the FlowSOM modality used in Spectre, carrying out clustering based on a self-organizing map (SOM) method, while Seurat first ran PCAs on the overall dataset, next constructed a K-nearest neighbour (KNN) graph, similar to PhenoGraph, another single cell analytical tool, and then adopted the Louvain algorithm to group cells together [10,24]. Such differences might account for the discrepancies we observed when comparing the clustering outcomes from Seurat and Spectre. Detailed comparisons of these mathematical methodologies are not within the scope of the current work, but previous study has shown that FlowSOM and PhenoGraph were the top-performing unsupervised methods for mass cytometry data clustering analysis. The KNN graph model deployed in PhenoGraph excelled in its clustering precision, stability and robustness in identifying sub-clusters, relative to other approaches like flowMeans, DEPECHE and Xshift [52]. Since Seurat shares the similar KNN model to PhenoGraph, it is reasonable to expect it could also show robust capacity in more generic cytometric data analysis. This warrants more systematic comparisons in future work. On the other hand, differences in results from Seurat and Spectre highlight that utilising both methods in tandem will provide a more complete understanding of complex datasets.

In addition to its distinct mathematical nature, as one of the cutting-edge end-to-end analytical tools for single cell data, Seurat has already been widely used in various research and clinical settings and is vigorously maintained and supported by its broad user community. This contributes to its easy accessibility and high user-friendliness and might reduce the coding burden as well, as users, especially those with previous experience in scRNA-seq, would not need to learn a completely new package or coding language for analysis. Moreover, application of Seurat potentially opens more possibilities for cytometric data analysis. As a popular scRNA-seq analysis package, Seurat could also act as a wrapper with favourable interoperability around a wide range of complementary packages or plugins originally developed for scRNA-seq analysis, such as LIGER and Harmony for data integration [29,53]. They might also be applicable to cytometric data, such as for batch correction, and could provide novel possibilities for high dimensional data analyses once validated. Thus, adapting Seurat offers a single simple platform to analyse, compare and cross-validate protein and RNA, and even potentially other multi-omic single cell data.

Previously, there were only few reports applying Seurat for protein level single cell analysis, such as for cellular indexing of transcriptomes and epitopes by sequencing (CITE-seq) [27] and CyTOF [54,55], and leveraging the rPCA integration method in Seurat for spectral cytometry analysis [35,56]. Recently, there has also been a similar attempt, adapting Scanpy, a Python-based scRNA-seq analysis package, to analyse mass cytometry data [57]. To our knowledge, the present work represents the first example of applying Seurat as a complete flow cytometric analysis workflow. Harnessing our newly developed 20-colour antibody panel and the Seurat-based analysis pipeline, we reported a unique T cell subset in CBMCs, characterised as CD8^+^CD45RA^+^CD27^+^CD161^+^ T cells. This subset partly overlapped with the previously described CD8^+^CD161^+^ T cells [41]. Previously, studies first discovered the differential (low/intermediate/high) levels of CD161 on CD8^+^ T cells in AB, which correlated with their various functional activities including cytokine production, proliferation, and lytic activity [58]. AB CD8^+^CD161^hi^ T cells were predominantly mucosal associated invariant T (MAIT) cells [59], while the CD8^+^CD161^int^ population represented a memory T cell subset which were enriched in the colonic lamina propria [41]. Consistent with this, our clustering analysis found a CD8^+^CD161^+^ population predominantly existing in AB (Fig. 3B), although with the current clustering setting, both Seurat and Spectre failed to further subdivide it into CD161^int^ and CD161^hi^ subsets. As for our newly identified CD8^+^CD45RA^+^CD27^+^CD161^+^ subset, considering its naïve phenotype, it is not surprising that it is almost negligible in AB.

The role of the CD8^+^CD161^+^ T cells in CB remains elusive. Developmentally, it was found that the CD8^+^CD161^hi^ T cells in CB might be the progenitor for post-natal MAIT cells [42,59,60]. Functionally, the CD161^hi^ subset produced IFN-γ and IL-17 [60,61], while the CD161^int^ subset, despite expressing markers like CD45RA and CCR7, still exhibited a preprogrammed transcriptomic profile reflective of their AB counterpart [41]. Our current clustering analysis could not further separate the population based on CD161 levels, but adjusting the clustering parameters could potentially help to differentiate them considering their intermediate to high CD161 expression (Fig. 4B). The naïve phenotype of this CB-enriched subset, based on the expression of CD45RA and CD27, is similar to previous reports [41,61], and we also confirmed its IFN-γ production. Previously, there were limited studies investigating the functional surface markers of CD8^+^CD161^+^ T cells, such as CCR6 [59]. Here, we characterised the CD8^+^CD45RA^+^CD27^+^CD161^+^ population as high in integrin β7 but low in CLA expression, implying preference to gut over skin homing. Thus, this CB population might represent the progenitors for AB CD8^+^CD161^int^ T cells which are enriched within the colon [41]. CD8^+^CD161^+^ T cells in AB are involved in the response to tissue-localized inflammation trigged by intracellular and viral pathogens [41,59], while their functional implications in CB remains elusive. Likewise, both our flow cytometry and scRNA-seq data suggested the pro-inflammatory and cytotoxic properties of CB CD8^+^CD45RA^+^CD27^+^CD161^+^ T cells. Additionally, as CD161 contributes to prenatal immune suppression [62], this subset might be involved in maintaining tolerance in the semi-allogenic context of pregnancy.

In summary, we have adapted Seurat, a widely used scRNA-seq analysis package, for high dimensional flow cytometric data analysis and showcased its performance through the identification of a unique CD8^+^CD45RA^+^CD27^+^CD161^+^ T cell population in CB. Such a pipeline presents a novel avenue for comprehensive analysis of high dimensional complex cytometric and multi-modal data, facilitating unbiased data-driven studies and discovery.

## Acknowledgements

We sincerely thank the Sydney Cytometry Core Research Facility for providing access to flow cytometric analysers. Part of the components within the figures were created with Servier Medical Art template, which are licensed under a Creative Commons Attribution 3.0 Unported License: https://smart.servier.com. This project was funded by the National Health and Medical Research Council (Australia, APP1104134) and the Norman Ernest Cummings Bequest. D.N. is a recipient of the Australian Government Research Training Program Scholarship (International). F.M-W. is supported by the International Society for the Advancement of Cytometry (ISAC), Marylou Ingram Scholars Program.

## Author Contributions

J.G.A.R. and D.N. together carried out all the analyses, participated in the project design and wrote the manuscript. B.S-N and G.V.P. performed the flow cytometry experiments. L.K., T.M.A., F.M-W., A.G.S., C.L.W. and J.T. provided intellectual input with the data analysis. P.H., N.J.C.K., and L.M. helped with the study design. R.N. designed the project, supervised the study, and wrote the manuscript. All authors reviewed and edited the manuscript.

## Figure Legends

**Supplementary Fig. 1**

**A.** Gating strategy to demultiplex and analyse the T cell subsets from CD3^+^CD4^+^ or CD3^+^CD8^+^ T cell compartments.

**B.** Scatter dot plots of the proportions of CD4^+^ and CD8^+^ T cells among the total lymphocytes (CD45^+^) in adult blood (AB) versus cord blood (CB).

N=5 per group and data are presented as mean ± s.e.m.

**Supplementary Fig. 2**

**(A-B).** Bar charts, with representative FACS gating, visualising the proportions of cells expressing CD49b^+^ **(A)** and FCER1A^+^ **(B)** from each cluster identified by Seurat.

**C.** Bar chart comparing the proportions of the clusters identified by Seurat (blue) and Spectre (red) within the total CD8^+^ T cell compartment. Clusters uniquely identified by one method are circled.

**D.** Uniform manifold approximation and projection (UMAP) plots visualizing the clustering results from Seurat (left) and Spectre (right). Cells that are differently clustered by Seurat and Spectre are circled. Sp2 from Spectre (i) is divided into Se1, Se2 and Se8 by Seurat (I). Se4 from Seurat (II) is split into Sp1 and Sp4 by Spectre (ii).

N=5 per group and data are presented as mean ± s.e.m.

**Supplementary Fig. 3**

**A.** Manual gating strategy to identify the 14 clusters obtained by Seurat based on the adult PBMC CD8^+^ T cell experiment.

**Supplementary Fig. 4**

**A.** Uniform manifold approximation and projection (UMAP) plots visualizing the clustering results from Seurat (left) and Spectre (right) based on the adult peripheral blood mononuclear cells (PBMCs) CD4^+^ T cell experiment. One colour represents one cluster.

**B.** Dot plots visualizing the clusters identified by Seurat (left) and Spectre (right) and their marker expression profiles. The size of the dot corresponds to the percentage of cells expressing the corresponding markers and the colour gradient reflects the average normalised expression of the corresponding markers.

**C.** Venn diagram comparing the clustering results from Seurat and Spectre. Both methods were set to generate 15 clusters and 12 out of 15 clusters could be identified by both methods, while Se2, Se8, and Se15 could only be identified by Seurat and Sp1, Sp8 and Sp15 could only be identified by Spectre.

**D.** Bar chart comparing the proportions per sample within the total CD4^+^ T cell compartments of the clusters identified by Seurat or retrieved by manual gating.

N=5 per group and data are presented as mean ± s.e.m.

**Supplementary Fig. 5**

(**A-B**). Scatter dot plots of the proportions of CD4^+^ (**A**) and CD8^+^ (**B**) T cells expressing different surface and intracellular markers in adult blood (AB) versus cord blood (CB).

**(C-D).** Scatter dot plots of the proportions of CD4^+^ (**C**) and CD8^+^ (**D**) T cells expressing IFN-γ, IL-4, IL-5, IL-10, and IL-13 in adult blood (AB) versus cord blood (CB).

(**E-F**). Clustering results retrieved by Spectre for CD8^+^ T cells based on the combined AB and CB experiment. **E.** Uniform manifold approximation and projection (UMAP) plot visualizing clustering results from Spectre. **F.** Dot plot visualizing the clusters identified by Spectre and their marker expression profiles. The size of the dot corresponds to the percentage of cells expressing the corresponding markers and the colour gradient reflects the average normalised expression of the corresponding markers.

N=5 per group and data are presented as mean ± s.e.m., with * p<0.05 and ** p<0.01 by unpaired Mann-Whitney t-test.

**Supplementary Fig. 6**

**A.** Gating strategy of adult blood (AB) CD8^+^ T cells to identify the 13 clusters obtained by Seurat.

**B.** Bar chart comparing the proportions within CD8^+^ T cell compartments of the clusters identified by Seurat or gated out by manual analysis in AB.

N=5 per group and data are presented as mean ± s.e.m.

**Supplementary Fig. 7**

**A.** Gating strategy of cord blood (CB) CD8^+^ T cells to identify the 13 clusters obtained by Seurat.

**B.** Bar chart comparing the proportions within CD8^+^ T cell compartments of the clusters identified by Seurat or gated out by manual analysis in CB.

N=5 per group and data are presented as mean ± s.e.m.

**Supplementary Fig. 8**

**A.** Bar chart comparing the proportions within CD8^+^ T cell compartment of the clusters identified by Seurat in adult blood (AB) versus cord blood (CB).

N=5 per group and data are presented as mean, with * p<0.05 and ** p<0.01 by unpaired Mann-Whitney t-test.

**Supplementary Fig. 9**

**A.** The fraction of four clusters of naïve CD8^+^ T cell subsets in foetal spleen, cord blood and adult samples. Each colour denotes one cluster and the number at the bottom shows the numbers of cells sequenced from each sample.

