## Supplementary Tables for "A unique human cord blood CD8^+^CD45RA^+^CD27^+^CD161^+^ T cell subset identified by flow cytometric data analysis using Seurat"

**Supplementary Table 1.** Fluorophores and their combinations used for the anti-CD45 barcoding system.

| **Fluorophore**  **Sample ID** | **Pacific Blue** | **PerCP** | **Spark Blue** |
| --- | --- | --- | --- |
| **1** | **+** | **-** | **-** |
| **2** | **+** | **+** | **-** |
| **3** | **-** | **+** | **-** |
| **4** | **-** | **+** | **+** |
| **5** | **-** | **-** | **+** |

**Supplementary Table 2.** List of antibodies used in this study.

| Reagent | Clone | Source | Identifier |
| --- | --- | --- | --- |
| Antibodies |  |  |  |
| Anti-human CD45 Pacific Blue | HI30 | Biolegend | Cat# 304029; RRID: AB_2174123 |
| Anti-human CD45 PerCP | HI30 | Biolegend | Cat# 304026; RRID AB_893337 |
| Anti-human CD45 Spark Blue | 2D1 | Biolegend | Cat# 368558; RRID AB_2904402 |
| Anti-human CD3 BUV496 | UCHT1 | Becton Dickinson | Cat# 612940; RRID: AB_2870222 |
| Anti-human CD4 BUV805 | SK3 (Leu3a) | Becton Dickinson | Cat# 612887; RRID: AB_2870176 |
| Anti-human CD8 BUV805 | SK1 | Becton Dickinson | Cat# 612889 RRID: AB_2833078 |
| Anti-human CD45RA BUV395 | 5H9 | Becton Dickinson | Cat# 740315; RRID: AB_2740052 |
| Anti-human CD27 BUV737 | O323 | Becton Dickinson | Cat# 751681; RRID: AB_2875667 |
| Anti-human Integrin β7 (β7) BB700 | FIB504 | Becton Dickinson | Cat# 746193; RRID: AB_2743541 |
| Anti-human CD161 PE-Cy5 | DX12 | Becton Dickinson | Cat# 551138; RRID: AB_394068 |
| Anti-human PSGL-1 (CLA) BB630/P | HECA-452 | Becton Dickinson | Cat# 624294 |
| Anti-human CXCR-5 BUV563 | RF8B2 | Becton Dickinson | Cat# 741316; RRID: AB_2870835 |
| Anti-human LAG-3 APC/R700 | T47-530 | Becton Dickinson | Cat# 565774; RRID: AB_2744329 |
| Anti-human FcERI (FCER1A) BV510 | AER-37 (CRA-1) | Becton Dickinson | Cat# 334626; RRID: AB_2564291 |
| Anti-human CD49b BV650 | AK-7 | Becton Dickinson | Cat# 560672 |
| Anti-human CD137 BV786 | 4B4-1 | Becton Dickinson | Cat# 741000; RRID: AB_2740623 |
| Anti-human CRTH2 BV711 | BM16 | Becton Dickinson | Cat# 740813; RRID: AB_2740474 |
| Anti-human CD40L BV605 | 24-31 | Biolegend | Cat# 310826; RRID: AB_2563832 |
| Anti-human T-box protein 21 (T-bet) BV421 | O4-46 | Becton Dickinson | Cat# 563318; RRID: AB_2687543 |
| Anti-human GATA-3 PE-Cy7 | L50-823 | Becton Dickinson | Cat# 560405; RRID: AB_1645544 |
| Anti-human BCL-6 APC | REA373 | Miltenyi Biotech | Cat# 130-121-997; RRID: AB_2801823 |
| Anti-human Forkhead box protein P3 (FoxP3) AF488 | 259D | Biolegend | Cat# 320212; RRID: AB_430887 |
| Anti-human Nur77 PE | 12.14 | eBioscience | Cat# 12-5965-82; RRID: AB_1257209 |
| LIVE/DEAD™ Fixable Near-IR Dead Cell Stain |  | Thermofisher | Cat# L34976 |
| Anti-human IL-13 APC | JES10-5A2 | Becton Dickinson | Cat# 561162; RRID: AB_10642586 |
| Anti-human IFN-γ BV421 | B27 | Becton Dickinson | Cat# 562988; RRID: AB_2737934 |
| Anti-human IL-5 FITC | REA1025 | Miltenyi Biotech | Cat# 130-117-203; RRID: AB_2727871 |
| Anti-human IL-10 PE | JES3-19F1 | Becton Dickinson | Cat# 559330; RRID: AB_397227 |
| Anti-human IL-4 PE-Cy7 | 8D4-8 | Becton Dickinson | Cat# 560672; RRID: AB_1727547 |

**Supplementary Table 3.** Classification, feature marker expression and functional implications for adult blood (AB) T cell clusters identified by Seurat.

| Cluster | Class* | Markers | Potential Properties |
| --- | --- | --- | --- |
| Se1 | Naïve | CD45RA^hi^ CD27^hi^ β7^hi^ | High gut homing potential. |
| Se2 | Naïve | CD45RA^hi^ CD27^hi^ β7^int^ | Intermediate gut homing potential. |
| Se3 | - | CD45RA^int^ CD27^low^ | Terminally differentiated and begin to re-express CD45RA. |
| Se4 | CM | CD45RA^low^ CD27^hi^ β7^hi^ CXCR-5^int^ | Follicular T cells with high gut homing potential. |
| Se5 | CM | CD45RA^low^ CD27^hi^ CLA^hi^ | Skin homing. |
| Se6 | - | CD45RA^int^ CD27^low^ CD161^int^ | Terminally differentiated cytotoxic phenotype and begin to re-express CD45RA. |
| Se7 | CM | CD45RA^low^ CD27^hi^ β7^hi^ CD161^hi^ | Cytotoxic with high gut homing potential |
| Se8 | Naïve | CD45RA^hi^ CD27^hi^ β7^low^ | Low gut homing potential. |
| Se9 | - | CD45RA^hi^ CD27^int^ β7^int^ CD161^int^ | Cytotoxic with intermediate gut homing potential.  Higher propensity for differentiation. |
| Se10 | - | CD45RA^int^ CD27^int^. | Transitioning phenotype from naïve to fully differentiated memory. |
| Se11 | EM | CD45RA^low^ CD27^low^ β7^low^ | Low gut homing potential. |
| Se12 | EMRA | CD45RA^hi^ CD27^low^ β7^int^ | Intermediate gut homing potential. |
| Se13 | EM | CD45RA^low^ CD27^low^ β7^hi^ | High gut homing potential. |
| Se14 | - | CD45RA^int^ CD27^int^ LAG-3^hi^ CXCR-5^hi^ | Recently activated follicular T cells. |

* T cells are classified into 4 subsets based on the expression of CD45RA and CD27: CD45RA^+^CD27^+^ (naïve); CD45RA^-^CD27^+^ (central memory; CM); CD45RA^-^CD27^-^ (effector memory; EM) and CD45RA^+^CD27^-^ (effector memory cells that re-express CD45RA; EMRA).

**Supplementary Table 4.** Classification, feature marker expression and functional implications for AB and CB T cell clusters identified by Seurat.

| Cluster | Class | Markers | Potential Properties |
| --- | --- | --- | --- |
| 1 | Naïve | CD45RA^hi^ CD27^hi^ β7^low^ | Low gut homing potential. |
| 2 | Naïve | CD45RA^hi^ CD27^hi^ β7^hi^ | High gut homing potential. |
| 3 | CM | CD45RA^int^ CD27^hi^ β7^hi^ | High gut homing potential. |
| 4 | CM | CD45RA^int^ CD27^hi^ CLA^hi^ | Skin homing. |
| 5 | EMRA | CD45RA^hi^ CD27^low^ β7^low^ | Low gut homing potential. |
| 6 | - | CD45RA^int^ CD27^low^ CD161^int^ | Terminally differentiated cytotoxic phenotype and begin to re-express CD45RA. |
| 7 | CM | CD45RA^low^ CD27^hi^ β7^hi^ CXCR-5^hi^ | Follicular T cells with high gut homing potential. |
| 8 | Naïve | CD45RA^hi^ CD27^hi^ β7^hi^ CD161^int^ | Cytotoxic with high gut homing potential. |
| 9 | - | CD45RA^int^ CD27^int^ β7^int^ CD161^hi^ | Transitory phenotype from naïve to fully differentiated memory.  Cytotoxic with intermediate gut homing potential. |
| 10 | - | CD45RA^int^ CD27^int^ | Transitory phenotype from naïve to fully differentiated memory. |
| 11 | EM | CD45RA^low^ CD27^low^ β7^hi^ | High gut homing potential |
| 12 | EMRA | CD45RA^hi^ CD27^low^ β7^int^ CD161^int^ | Cytotoxic with intermediate gut homing potential. |
| 13 | - | CD45RA^int^ CD27^int^ LAG-3^hi^ CXCR-5^hi^ | Recently activated follicular T cells |

* T cells are classified into 4 subsets based on the expression of CD45RA and CD27: CD45RA^+^CD27^+^ (naïve); CD45RA^-^CD27^+^ (central memory; CM); CD45RA^-^CD27^-^ (effector memory; EM) and CD45RA^+^CD27^-^ (effector memory cells that re-express CD45RA; EMRA).
