## Supplementary Figures for "A unique human cord blood CD8^+^CD45RA^+^CD27^+^CD161^+^ T cell subset identified by flow cytometric data analysis using Seurat"

### Supplementary Fig. 1

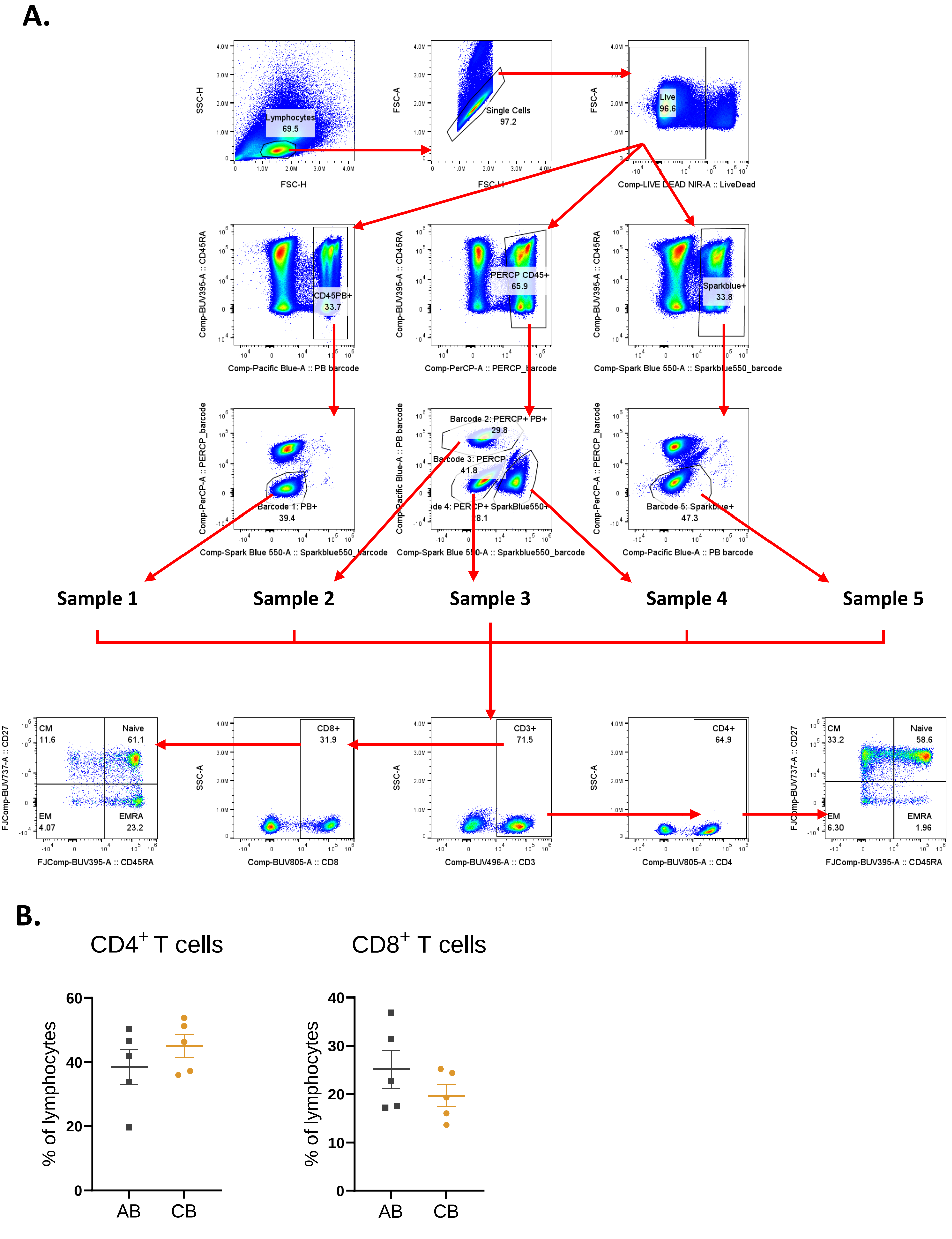

### Supplementary Fig. 2

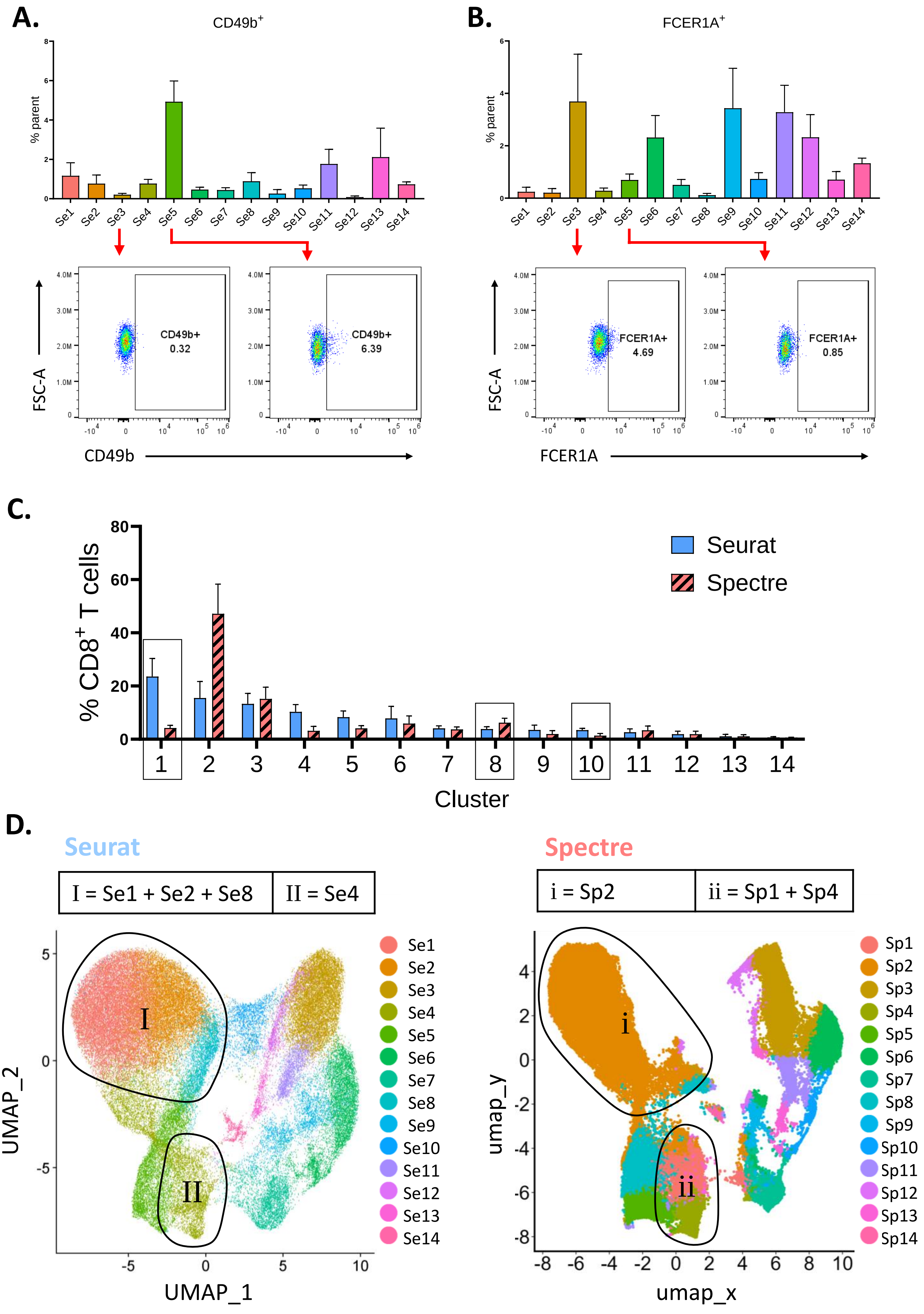

### Supplementary Fig. 3

**A.**

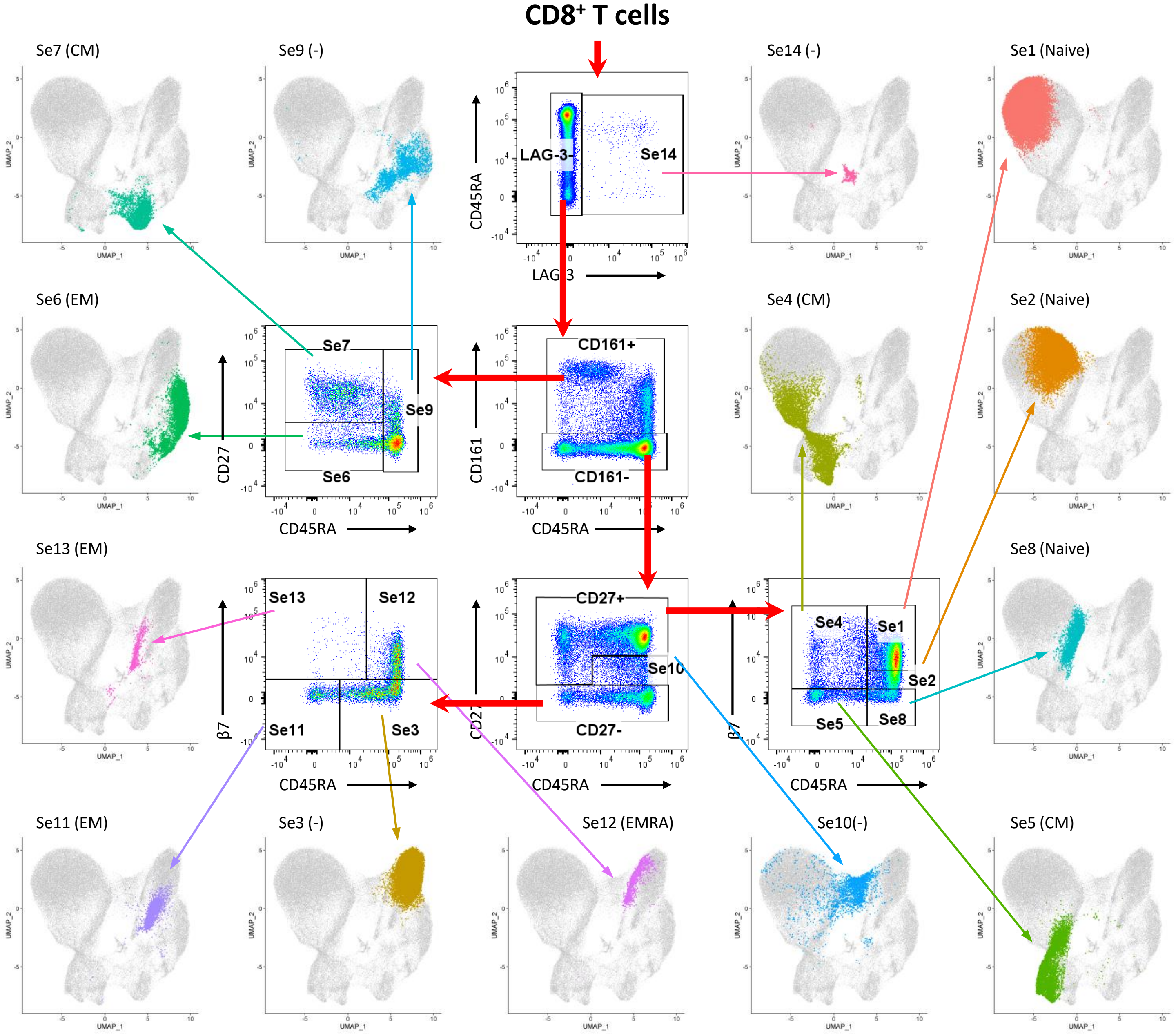

### Supplementary Fig. 4

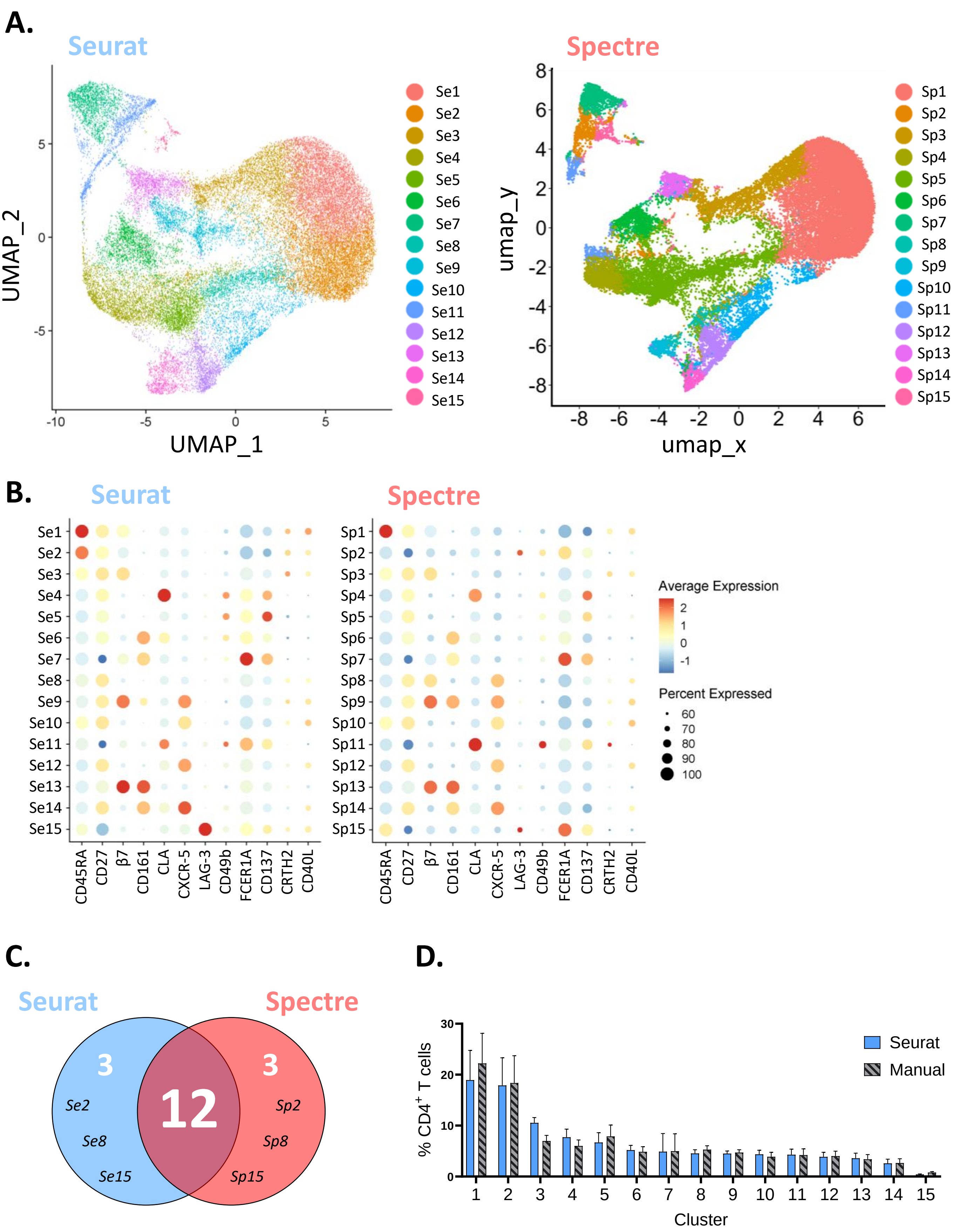

Supplementary Fig. 5

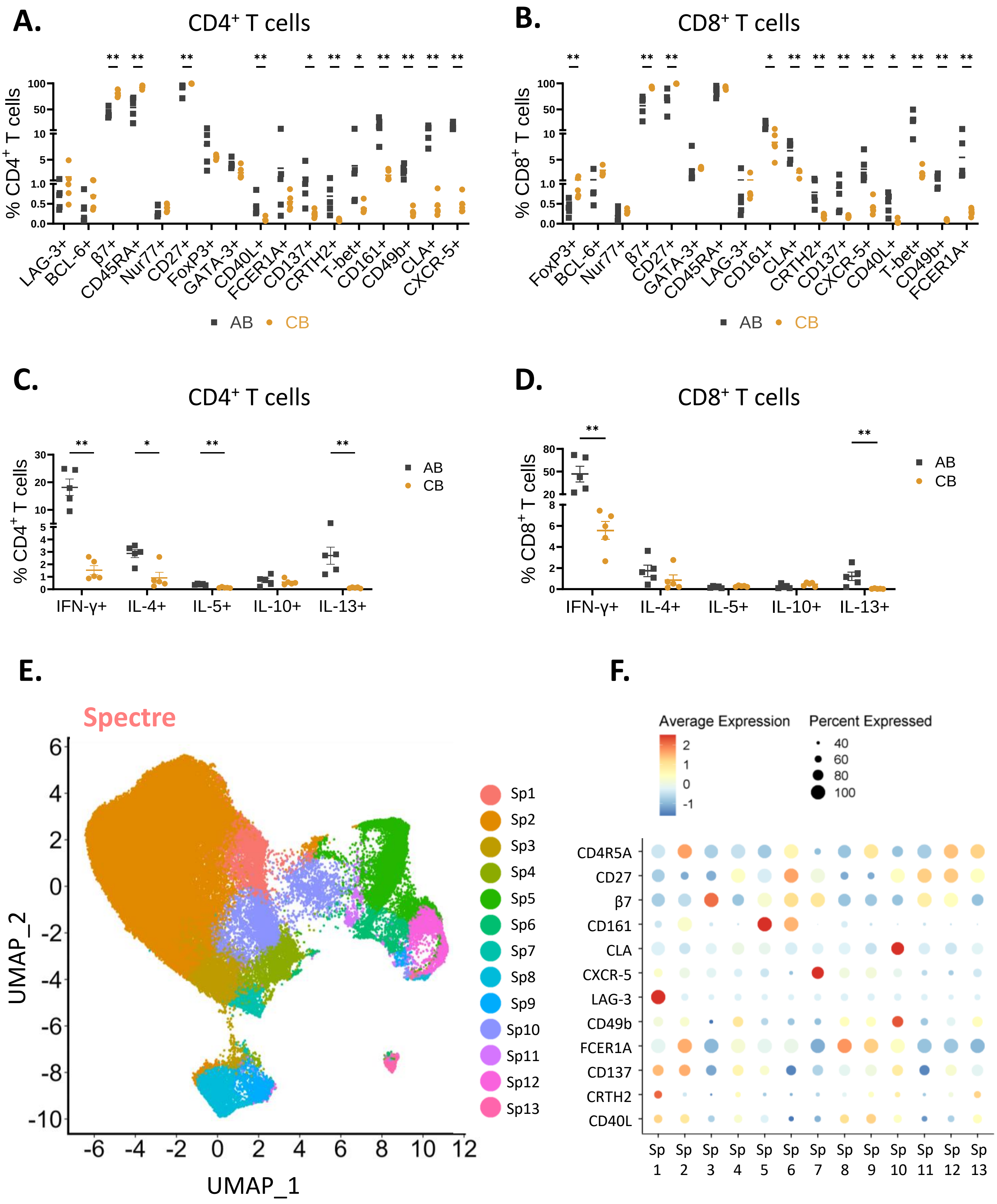

Supplementary Fig. 6

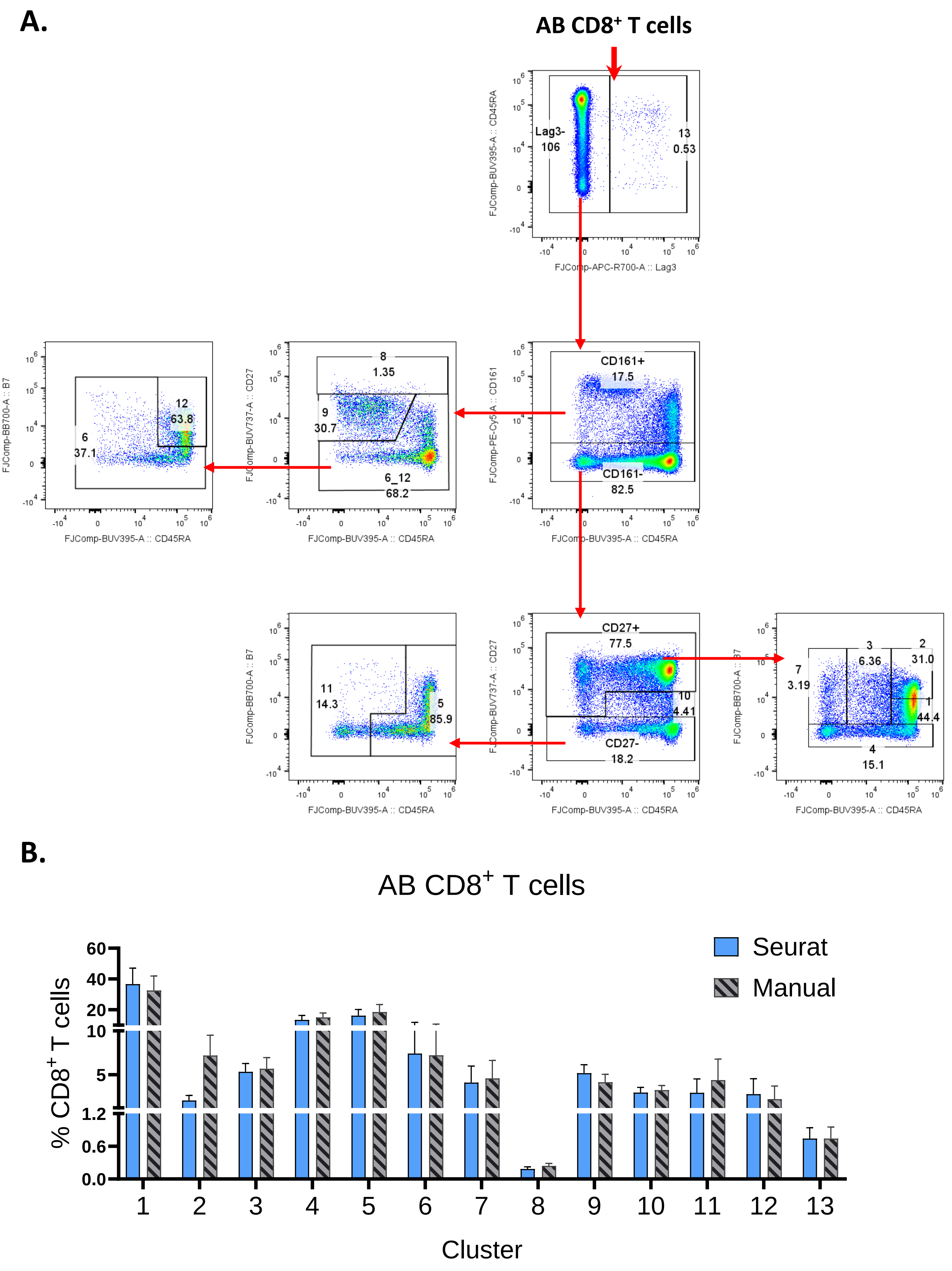

Supplementary Fig. 7

**A.**

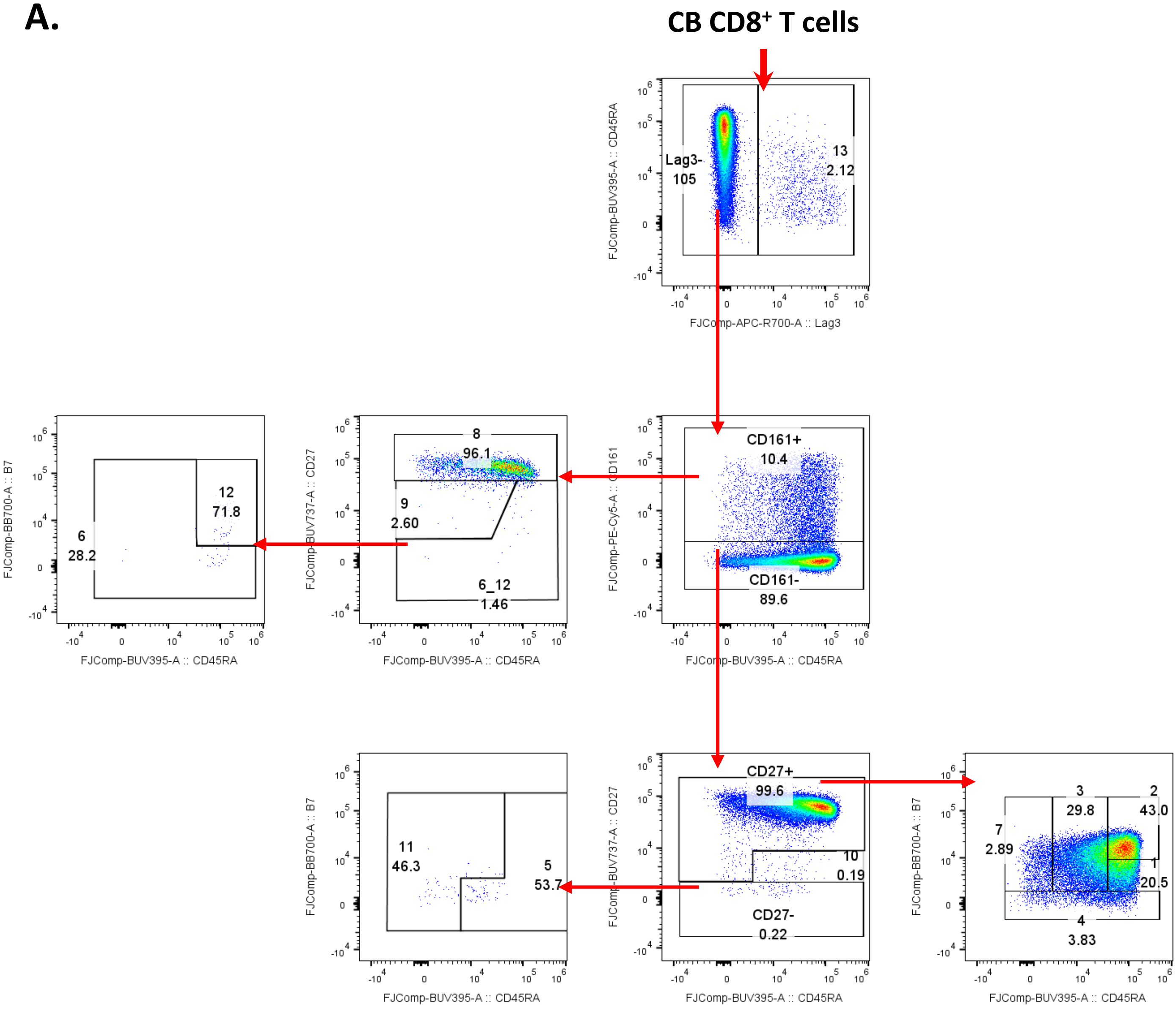


**B.**

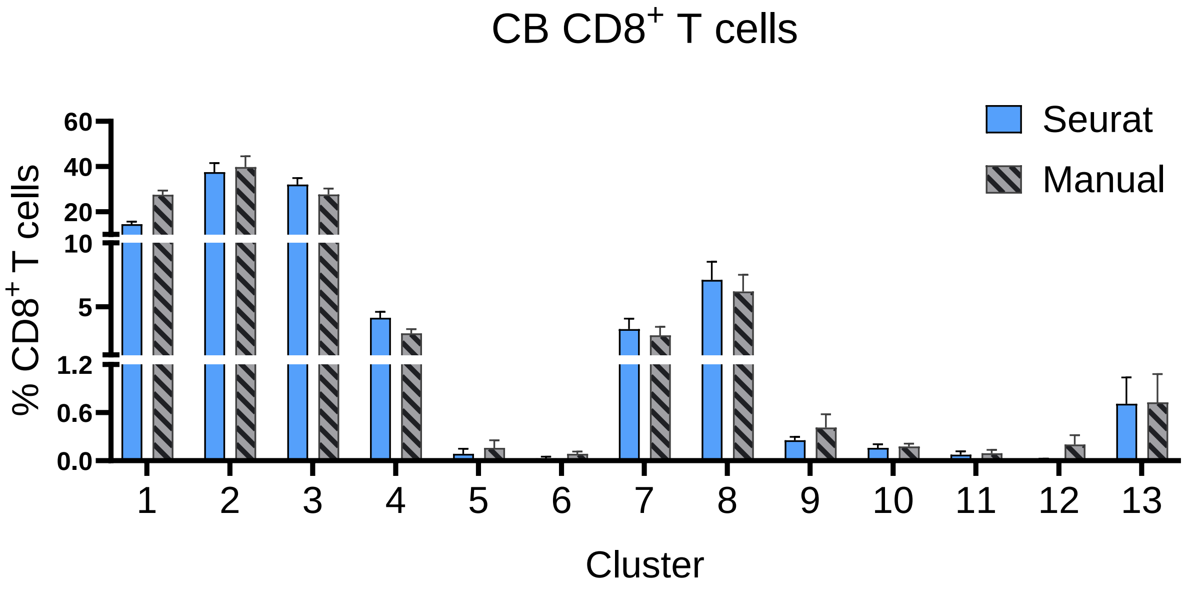

Supplementary Fig. 8

**A.**

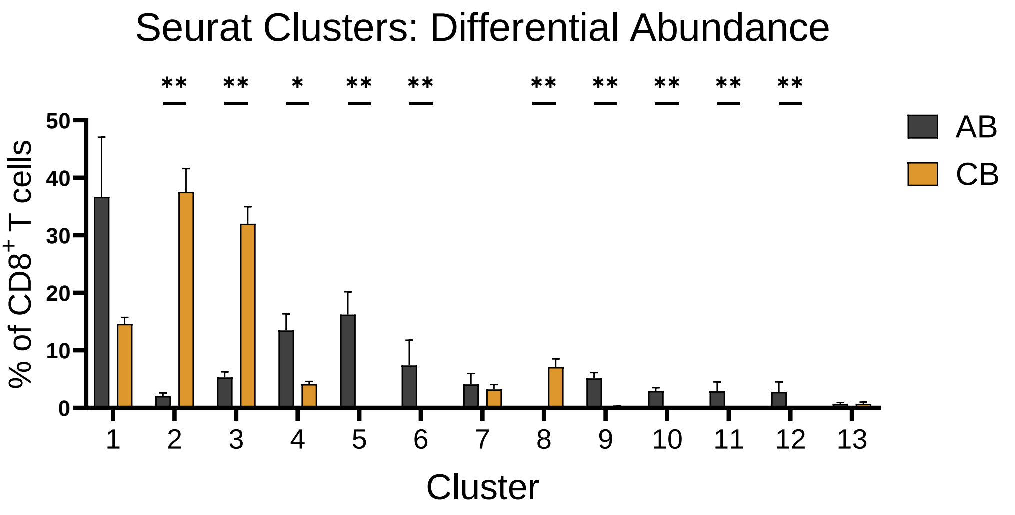

Supplementary Fig. 9

**A.**

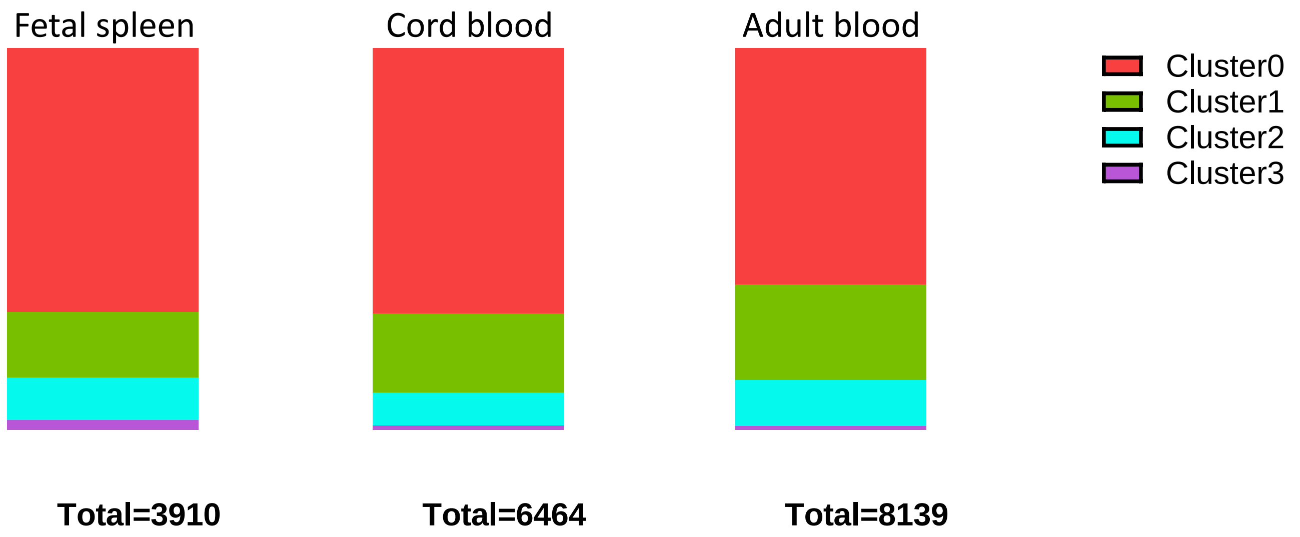
